# SirT7 auto-ADP-ribosylation regulates glucose starvation response through macroH2A1.1

**DOI:** 10.1101/719559

**Authors:** Nicolás G. Simonet, Joshua K. Thackray, Berta N. Vazquez, Alessandro Ianni, Maria Espinosa-Alcantud, Julia Morales-Sanfrutos, Sarah Hurtado-Bagès, Eduard Sabidó, Marcus Buschbeck, Jay Tischfield, Carolina de la Torre, Manel Esteller, Thomas Braun, Mireia Olivella, Lourdes Serrano, Alejandro Vaquero

## Abstract

Sirtuins are key players in the response to oxidative, metabolic and genotoxic stress, and are involved in genome stability, metabolic homeostasis and aging. Originally described as NAD^+^-dependent deacetylases, some sirtuins are also characterized by a poorly understood mono-ADP-ribosyltransferase (mADPRT) activity. Here we report that the deacetylase SirT7 is a dual sirtuin as it also features auto-mADPRT activity. Molecular and structural evidence suggests that this novel activity occurs at a second previously undefined active site that is physically separated in another domain. Specific abrogation of this activity alters SirT7 chromatin distribution, suggesting a role for this modification in SirT7 chromatin binding specificity. We uncover an epigenetic pathway by which ADP-ribosyl-SirT7 is recognized by the ADP-ribose reader macroH2A1.1, a histone variant involved in chromatin organization, metabolism and differentiation. Glucose starvation (GS) boosts this interaction and promotes SirT7 relocalization to intergenic regions in a macroH2A1-dependent manner. Both SirT7 activities are in turn required to promote GS-dependent enrichment of macroH2A1 in a subset of nearby genes, which results in their specific up- or downregulation. Consistently, the expression changes of these genes associated to calorie restriction (CR) or aging are abrogated in *SirT7*^-/-^ mice, reinforcing the link between Sirtuins, CR and aging. Our work provides a novel perspective about sirtuin duality and suggests a key role for SirT7/macroH2A1.1 axis in mammalian glucose homeostasis, calorie restriction signaling and aging.

---

The members of the Sir2 family, also known as sirtuins, play a key role in the response to different types of stress, including metabolic, oxidative and genotoxic stress. Thus, sirtuins are at the crossroads of many pathways that control genome stability, metabolic homeostasis, cell cycle control and apoptosis (Bosch-Presegue and Vaquero, 2014). The family members originated in the prokaryotes, where they participated in the metabolism of NAD^+^ (Frye, 2000). During evolution, sirtuins have expanded and diversified, acquiring a key role in the crosstalk between chromatin and environment. Reflecting this, four of the seven mammalian sirtuins (SirT1, 2, 6, 7) have been linked to the control of genome stability at different levels (Martínez-Redondo and Vaquero, 2013). One of the most intriguing features of sirtuin biology is that they catalyze two NAD^+^-dependent enzymatic activities: NAD^+^-dependent deacetylation and mono-ADP ribosylation (mADPR), which involves the reversible transfer of one ADP-ribose moiety from the coenzyme NAD^+^ to specific protein residues (Bheda et al., 2016). The former seems to be more prominent among mammalian sirtuins, since almost all of them are NAD^+^-dependent protein deacetylases, whose activity can, in some cases, target other single acyl groups such as butyryl or succinyl, or even long-chain fatty acyl groups (Jiang et al., 2003). The relevance of this duality goes beyond sirtuins, as we know of just a handful of enzymes, most of them prokaryotic, that combine two activities in a single protein (Caban and Ginsburg, 1976; Laporte et al., 1989; Nureki et al., 1998).

By far the best studied ADP-ribosyltransferase is PARP1, which is responsible for up to about 90% of the cellular poly-ADP-ribosylation (pADPR) generated by DNA damage, and is a key pharmacological target for the treatment of DNA repair-deficient cancers (Barkauskaite et al., 2013; Brown et al., 2016). In contrast to the relatively well studied pADPR, the biological relevance of mADPR is more limited. mADPR has been shown to regulate a wide range of functions, from cellular stress response upon DNA damage (Martello et al., 2016) or metabolic stress (Aguilera-Gomez et al., 2016), to development of infectious diseases and cancers (Bütepage et al., 2015; Ling et al., 2017; Scarpa et al., 2013). Reflecting their functional relevance and complexity, there are different classes of protein-binding modules (readers) that recognize ADP-ribosylation, but also enzymes involved in the removal of ADP-ribose called ADP-ribosylhydrolases (erasers) (Hottiger, 2015){Hottiger:2015es}. Among the readers, macrodomains are an ancient group of protein domains formed by 130-190 residues present from viruses to humans that bind MAR, PAR and NAD^+^-derived metabolites (Rack et al., 2016). In humans, at least 12 proteins contain macrodomains belonging to four of the six known classes of macrodomains. One of the best studied macrodomain-containing proteins is the histone H2A variant macroH2A (mH2A) (Sun and Bernstein, 2019). Of the three mH2A isoforms (mH2A1.1, 1.2 and 2), only mH2A1.1 has the ability to bind ADPR (Kustatscher et al., 2005). Moreover, mH2A1.1 can also bind free ADP-ribose molecules and the product of sirtuin deacetylation, O-acetyl-ADP-ribose (Pazienza et al., 2014), which suggests that mH2A1.1 may play a role as an energetic/metabolic sensor and could be functionally linked to the stress-response role of sirtuins. Interestingly, mH2A1.1 has been implicated in gene repression and activation (Chen et al., 2014; Gamble et al., 2010), in the latter case especially after signal-induced gene activation through PARP1 inactivation (Ouararhni et al., 2006).

Sirtuin-mediated ADP-ribosylation was described in yeast Sir2p before the discovery of the main deacetylation activity (Frye, 1999), and it was originally hypothesized that all sirtuins could potentially harbor mADPRT activity (Tanny et al 1999), although this activity is, in most cases, either orders of magnitudes slower than deacetylation or simply undetectable in *in vitro* studies (Fahie et al., 2009). Among mammalian sirtuins (SirT1 to SirT7), only two —reported to be weak deacetylase enzymes *in vitro* (Lin et al., 2012)— are known to ADP-ribosylate other protein targets. SirT4-dependent ADP-ribosylation of glutamate dehydrogenase (GDH) represses its enzymatic activity by limiting the metabolism of glutamate and glutamine to produce ATP (Haigis et al., 2006). SirT6 can mediate the ADP-ribosylation of two targets: PARP1 under oxidative stress, thereby inducing DSB repair after DNA damage (Mao et al., 2011); and KAP1, to promote LINE-1 silencing and genomic stability (Van Meter et al., 2014). SirT6 was also found to auto-ADP-ribosylate itself (Liszt et al., 2005a), although this activity has not been characterized. *In vitro* kinetic analysis suggests that the deacetylase activity of SirT6 with an acetylated H3K9 peptide is about 1000 times weaker than that of other sirtuins (Pan et al., 2011). Both enzymatic activities in SirT6 seem to involve specific residues within the same catalytic domain, since specific mutations in this domain can selectively abrogate either mADPRT or deacetylase without altering the other activity (Liszt et al., 2005b; Mao et al., 2011). The coexistence of the two activities in the same protein is uncommon and raises many questions about the nature of this duality, the crosstalk between the two activities, and their specific contribution to the physiological role of sirtuins.

SirT7 is the closest sirtuin to SirT6 and probably one of the least studied. Its deficiency has been linked to genome instability and several pathologies, including cancer and aging (Barber et al., 2012; Vazquez et al., 2016). SirT7 is involved in active rDNA transcription (Ford et al., 2006), DNA repair (Vazquez et al., 2016), metabolic homeostasis (Ryu et al., 2014; Yoshizawa et al., 2014), heterochromatin silencing and active transcriptional elongation through deacetylation of histone H3 lysine 18 (H3K18ac) and lysine 36 (H3K36ac), respectively (Barber et al., 2012; Wang et al., 2019). Recently, a direct link between SirT7, nuclear lamina and LINE-1 has been described, suggesting a role for SirT7 in genome organization (Vazquez et al., 2019). SirT7 knockout mice show severe metabolic defects, including resistance to high-fat diet-induced fatty liver, obesity, glucose intolerance and diminished levels of hepatic triglycerides and white adipocyte tissue (Fang et al., 2017; Yoshizawa et al., 2014). Additionally, these mice exhibit developmental defects and aging-related phenotypes such as perinatal lethality, degenerative heart hypertrophy, osteopenia, kyphosis, and they die significantly younger than their wild type counterparts (Fukuda et al., 2018; Vakhrusheva et al., 2008; Vazquez et al., 2016). While there is a large body of evidence on sirtuin function and longevity (Imai and Guarente, 2016), the mechanism by which SirT7 controls aging and the relationship between caloric restriction response and gene expression programs have not yet been adequately addressed.

Here we describe how SirT7 harbors an auto-mADPRT activity at an alternative catalytic site in addition to its primary catalytic site, which is responsible for the deacetylation reaction. Auto-mADPRT of SirT7 confers its binding capacity on specific chromatin regions through the histone variant mH2A1.1, an interaction that is boosted under glucose starvation (GS), which promotes accumulation of mH2A1 in a SirT7-dependent manner. To advance our understanding of SirT7 function under energy-stress conditions, we sought evidence of the key role of the SirT7/mH2A axis in calorie-restricted and aged mice. This axis regulates epigenetically the expression of genes involved in differentiation, proliferation, metabolism and aging pathways. Our functional studies suggest a direct role for SirT7 in the control of the stress response through this newly discovered enzymatic activity.

## Results

### SirT7 is a mono-ADP-ribosyltransferase

Several lines of evidence suggest that SirT7 may also harbor mADPRT activity. SirT7 and SirT6 are close homologues with a common origin in early eukaryotes. They consistently show a high level (42%) of sequence identity and structure in their catalytic core domain. In fact, SirT7 shares a unique feature with SirT6 in the catalytic domain that is different from other mammalian sirtuins (Fig. 1a). Whereas there is a large helix bundle in the NAD^+^-binding Rossmann fold in the catalytic region of most sirtuins that connects this domain with a small Zn-binding domain, this structure is missing from SirT6 and SirT7. This feature was thought to considerably reduce the flexibility of the structure, causing the low deacetylase activity detected in SirT6 (Pan et al., 2011). Consistent with this, the *in vitro* catalytic activity of SirT7 with H3K18Ac is considerably weaker than that of other sirtuins such as SirT1 or SirT2 (Wang et al., 2019). Considering all this evidence, we aimed to determine whether SirT7 was also a dual sirtuin and if it harbored an ADPRT activity. Our first approach was to employ an *in vitro* ADP-ribosylation assay, using [^32^P]-labeled NAD^+^ to determine whether SirT7 was able to transfer [^32^P]-ADP-ribose to recombinant core histones. Surprisingly, the ADPRT reactions of SirT7 on recombinant and native histones was very weak, but we detected a very efficient incorporation of [^32^P] in SirT7, strongly suggesting that SirT7 is itself a major target of this activity (Fig.1b and Supplementary Fig.1a). The absence of a smear in the [^32^P]-signal indicated that, as with SirT6, SirT7 harbors mADPRT activity (Fig.1c). Surprisingly, this auto-ADPRT activity was not altered by mutations previously described as abrogating SirT6 ADP-ribosylation activity towards PARP1 and KAP1 (Mao et al., 2011; Van Meter et al., 2014). Mutation of SirT6 G60 to alanine, or its equivalent in SirT7 S115 did not abrogate this auto-ADPRT activity. Moreover, mutation of SirT7 S111, equivalent to SirT6 S56, which was shown to eliminate both SirT6 activities, behaved similarly (Fig.1d). These observations led us to consider the possibility that sirtuin auto-mADPRT activity is not associated with the previously described active site.

**Fig.1.**
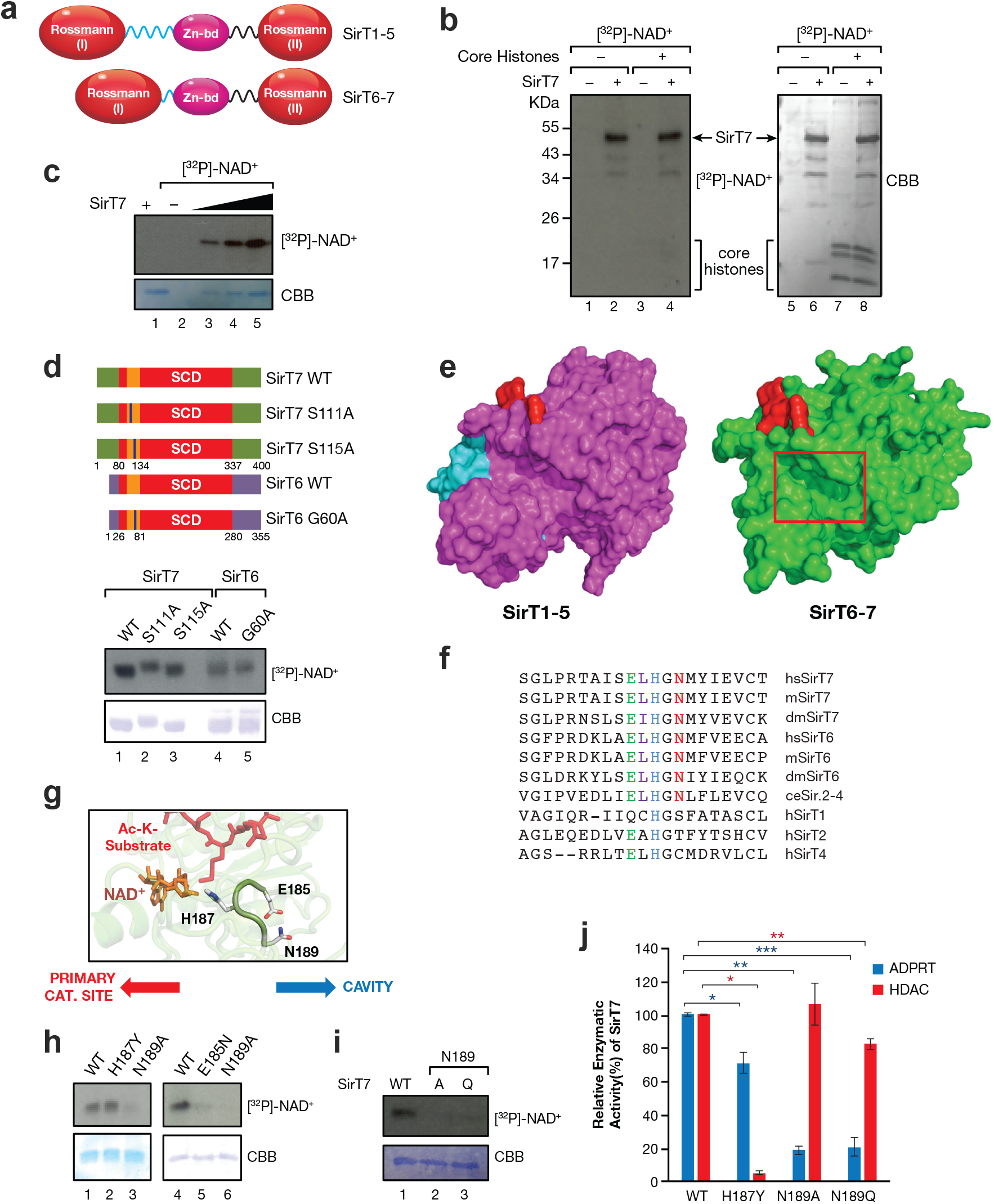
SirT7 harbors a mono-ADP-ribosyltransferase activity that is catalyzed by an alternative domain. **a**, Secondary structure of the conserved domain in mammalian sirtuins. Differences in α-helix length between SirT1-5 and SirT6-7 are shown in blue. **b**, *In vitro* ADPRT assay of purified SirT7 ± [^32^P]-NAD^+^ in the presence and absence of purified HeLa core histones. Autoradiography ([^32^P]-NAD^+^) and Coomassie-blue staining are shown. **c**, SirT7 titration in *in vitro* ADPRT assay performed as in b). **d**, Mutations in SirT7 and SirT6 primary sites do not abrogate SirT7 auto-ADPRTion. Upper panel, Schematic representation of SirT7 and SirT6 WT and mutants tested. Lower panel, *in vitro* ADP-ribosylation performed as in b) and c). **e**, Structural comparison of conserved domains of human SirT1-5 (in magenta) and SirT6-7 (in green). The main structural difference between SirT1-5 and SirT6-7 is an additional helix bundle subdomain in the latter. The primary catalytic site with a bound acetylated peptide is shown in red, while the α-helix bundle that is not present in SirT6 and SirT7, unlike in other sirtuins, is shown in cyan. Loss of this region induces the formation of a cavity in SirT6 and 7 (red square). **f**, Alignment of the sequence present in the cavity in the SirT6 and 7 lineage and the corresponding sequence in hSirT1, 2 and 4. Residues E185, H187 and N189 are shown in green, blue and red, respectively. **g**, Molecular model of the SirT7 catalytic domain based on the crystal structure of SirT6 (PDB 5M6F, 1.87Å). The catalytic residue histidine 187 (H187) of SirT7 is oriented towards the NAD^+^ molecule (in orange) and the acetylated substrate (in red). Residues E185 and N189, proposed to be involved in SirT7 auto-mADPRT activity, are located in the same loop as H187 although pointing in the opposite direction. **h-i**, *In vitro* auto-ADP-ribosylation assay of SirT7 WT and indicated mutants. **j**, Comparison of deacetylation and ADP-ribosylation activity of the indicated proteins. In each case, the results are shown relative to WT SirT7 activity (100%). Quantifications are based on three experiments. A representative deacetylation experiment is shown in Supplementary Fig. 1f (*p*: * <0.05; ** <0.01; ***<0.005).

### SirT7 auto-mADPRT activity depends on the highly conserved residues, E185 and N189, present in the ELGHN motif

In order to define this site, we compared all seven sirtuins at the structural level. In contrast to SirT1-5, the lack of the alpha helix bundle (in cyan; Fig.1e, left) in SirT6 and SirT7 (Fig.1e, right) induced a reorganization in the center of the structure that created a big cavity (Fig.1e, red rectangle), located at the other side of the main catalytic site (in red). The cavity contained an ELGHN motif, conserved in the SirT6/SirT7 lineage, including the common SirT6/7 ortholog *C. elegans* Sir.2-4 (Fig.1f, Supplementary Fig.1b). SirT7 H187, present in the motif, is a key conserved residue among sirtuins involved in the recognition of acetylated substrate in the deacetylation activity. While H187 was oriented towards the NAD^+^ binding pocket and the main catalytic site, the two flanking residues, E185 and N189, faced in the opposite direction, towards the surface of the cavity (Fig.1g, Supplementary Fig.1c-d), forming a loop sustained by interaction of both residues through their side chain. We paid particular attention to these two conserved residues because of their possible involvement in the ADP-ribosylation reaction: E185 is the only residue in this motif that could initiate the reaction, whereas N189, the only residue in the motif conserved exclusively in the whole SirT6/7 lineage (Fig.1f), could act as the first acceptor of the ADP-ribosyl moiety. This possibility was confirmed with the finding that the E185N and N189A mutations both abrogated SirT7 auto-mADPRT activity (Fig.1h).

A highly conservative mutation to glutamine (N189Q) had the same effect, suggesting that the lack of activity of the N189 mutant was not due to an indirect structural effect (Fig.1i). The fact that N189Q is structurally very similar to the WT protein (Supplementary Fig.1e) but cannot be ADP-ribosylated, suggests that this residue may play a role as a first acceptor of ADP-ribosylation. N189 mutants were defective in ADPRT activity but still featured deacetylase activity, while the H187Y mutant, just two residues away, had the completely opposite pattern (Fig.1j, Supplementary Fig.1f). Considered together, these observations indicate that SirT7 auto-mADPRtion is catalyzed at a conserved alternative secondary active site located in a previously uncharacterized domain.

### N189-dependent auto-mADPRT involves several residues distributed around the SirT7 surface and regulates SirT7 genomic distribution

We also confirmed the conserved role of N189, since the equivalent mutation in SirT6 N135 also abrogated SirT6 mADPRT activity (Fig.2a). There were two further lines of evidence of the importance of N189: first, the N189 mutant was significantly less ADP-ribosylated *in vivo* than was WT SirT7 (Fig.2b); second, mass spectrometry (MS) analysis of SirT7 auto-mADPRTion identified eight ADP-ribosylated peptides distributed across the entire surface of the SirT7 protein (Fig.2c-d, Supplementary Fig.2a-b and Table S1). All but one of them were undetected in the N189 mutant (Fig.2d, in magenta), confirming a key role for N189 in SirT7 ADPRT. Together, these observations strongly suggest that SirT7 auto-mADPRTion is catalyzed at an alternative secondary active site.

**Fig.2.**
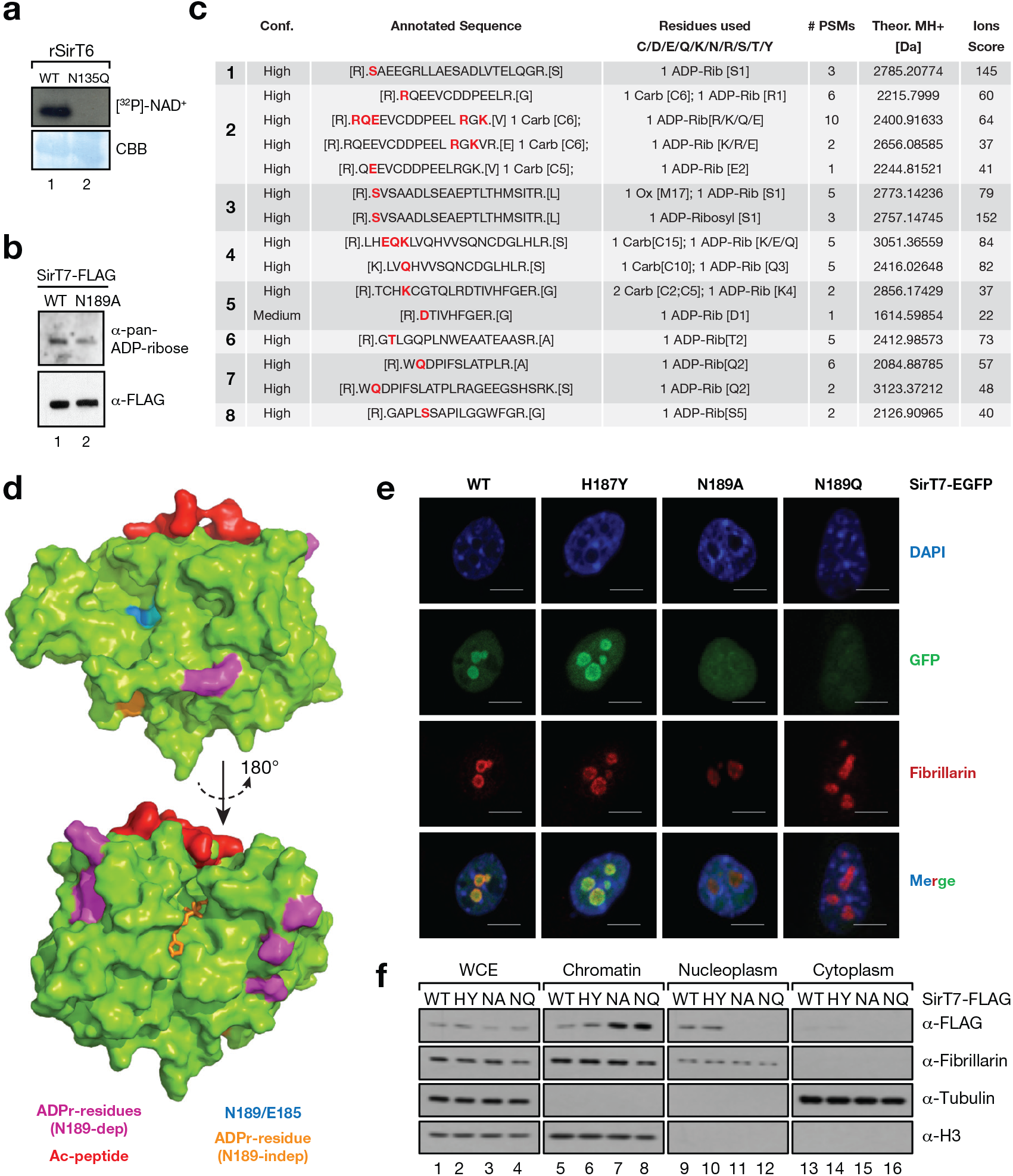
SirT7 N189-dependent auto-ADP-ribosylation regulates SirT7 distribution and chromatin-binding dynamics. **a**, *In vitro* auto-mAPDRT assay as before with bacterially expressed recombinant SirT6 WT or N135Q. **b**, ADP-ribosylation levels of SirT7 WT or N189A expressed in HEK293F cells monitored by far-western blot with anti-pan-ADP-ribose binding reagent. **c**, Analysis of SirT7 auto-mADPRTion identified by mass spectrometry. ADP-ribosylated peptides identified in SIRT7 WT or N189A were analyzed using HCD and EthcD fragmentation methods. Modification sites are highlighted in red. Further information is included in Table S1 and in the Methods. **d**, Structural model of the SirT7 catalytic domain indicating localization of the ADP-ribosylated peptides identified in c. The ADP-ribosylation N189-dependent (magenta) and independent (orange) residues, the N189/E185 cavity (blue) and the primary catalytic site bound to acetylated peptide (red) are shown. **e**, Immunofluorescence assay of the indicated GFP-tagged SIRT7 proteins. Fibrillarin was included as a nucleolar marker. Scale bar: 5 μm. **f**, Cellular distribution of SirT7 WT, H187Y (HH), N189A (NA) and N189Q (NQ) in whole-cell extract (WCE), cytoplasm, nucleoplasm and chromatin in NIH3T3 cells. Controls for the nuclear fraction (fibrillarin), cytoplasm (tubulin) and chromatin (histone H3) are also shown.

Taking advantage of our ability to specifically inhibit each of the two SirT7 enzymatic activities, we next studied the impact of these mutations on SirT7 distribution. Our first step was to study immunofluorescence (IF) signal of WT, H189Y, N189A and N189Q. We observed that while H189Y had an identical distribution to WT SirT7, whereby it was clearly enriched in the nucleolus, the patterns of localization of N189A and Q were radically different. Both N189 mutants showed a more dispersed pattern of distribution that resulted in a loss of nucleolar enrichment (Fig.2e). This was confirmed by biochemical fractionation of cells expressing SirT7-flagged mutants into cytoplasm, soluble nuclear fraction (nucleoplasm) and chromatin-insoluble fraction (chromatin). As in the IF studies, H187Y behaved similarly to WT and was present in similar levels in the nucleoplasm. By contrast, N189A-Q mutants were significantly enriched in the chromatin fractions and depleted in the nucleoplasm (Fig.2f). Overall, our results suggest that, in contrast to SirT7 deacetylase activity, mADPRT activity, and in particular N189, plays a key role in both SirT7 genomic localization and chromatin-binding dynamics.

### SirT7 interacts with the ADP-ribose binding protein mH2A1.1 upon glucose starvation

We tried to identify SirT7 binding partners to understand the functional link between SirT7 and ADP-ribosylation. We performed SirT7 affinity purification in HEK293F cells and identified potential SirT7 interactors by MS analysis. We identified the histone variant mH2A1 from among the candidates (Fig.3a and Supplementary Table S2). Remarkably, mH2A1.1, the only one of the three mH2A isoforms that specifically binds ADP-ribose through its macrodomain (Fig.3b) (Kustatscher et al., 2005) and interacted specifically with SirT7 (Fig.3c). We next examined whether the SirT7-mH2A1.1 interaction depends on SirT7 ADP-ribosylation by incubating bacterially expressed recombinant SirT7 (rSirT7) in the presence or absence of NAD^+^, and dialyzing the reaction to remove all remnant NAD^+^ molecules. We tested its ability to specifically pull down mH2A1.1 from HEK293F nuclear extracts under stringent conditions (Fig.3d). We found that only rSirT7 previously incubated with NAD^+^ was able to interact with WT mH2A1.1 under these conditions (Fig.3e). As evidence that SirT7 auto-mADPRT mediates this binding, no interaction was detected under these conditions when we used mH2A1.1 G224E (Fig.3e, lanes 5-6), a mutant unable to bind specifically to ADP-ribose (Kustatscher et al., 2005). This is compelling evidence that SirT7 auto-mADPRT mediates this binding.

**Fig.3.**
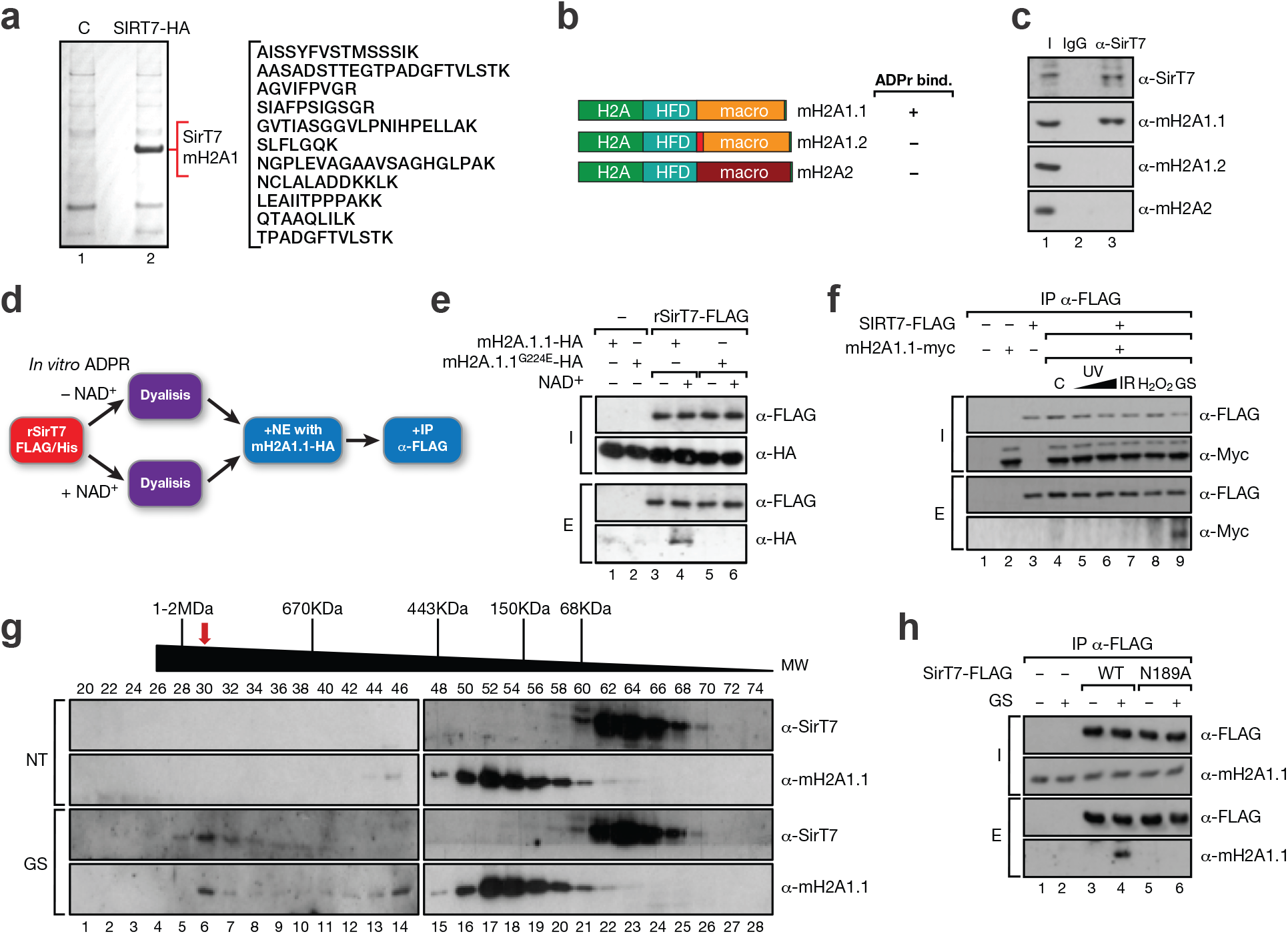
SirT7 interacts with mH2A1.1 upon GS in an ADP-ribosylation-dependent manner. **a**, Affinity purification of SirT7-binding factors identified the histone H2A variant mH2A1 by mass spectrometry analysis (Table S2). **b**, Schematic representation of the three mH2A isoforms, of which, only mH2A1.1 binds to ADP-ribose (Kustatscher et al., 2005). **c**, Endogenous SirT7 specifically immunoprecipitates mH2A1.1 in HEK293F cells. **d-e**, mH2A1.1 recognizes and binds ADP-ribosylated SirT7. Immunoprecipitation of bacterially expressed recombinant SirT7 pre-incubated ±NAD^+^ and added to nuclear extracts of HEK293F cells expressing mH2A1.1 WT or G224E, a mutant deficient in ADP-ribose binding. Inputs (I) and elutions (E) are shown. **f**, Interaction between SirT7 and mH2A1.1 under high-stringency conditions upon different types of stress. C: untreated; UV: ultra-violet, IR: 7-Gy ionizing irradiation; H_2_O_2_: oxidative stress; GS: glucose starvation. **g**, Superose 6 gel-filtration chromatography of nuclear endogenous proteins from HEK293F cells under normal cell growth (NT) or upon GS. Fraction numbers and approximate MWs are indicated. Western blot of SirT7 and mH2A1.1 are shown. **h**, High-stringency immunoprecipitation of endogenous mH2A1.1 by WT or N189A SirT7 in HEK293F cells treated under normal conditions or under GS.

The *in vivo* interaction between SirT7 and mH2A1.1 was specific, but weak. Considering the direct link between sirtuins and the stress response, we investigated whether the SirT7-mH2A1.1 interaction is boosted under specific stress conditions, such as UV or ionizing irradiation, oxidative stress (H_2_O_2_) or glucose starvation (GS). Under highly stringent conditions, there was a significant increase in the interaction between the two factors, specifically upon GS (Fig.3f), that was consistent with the established link between both proteins and glucose homeostasis (Pazienza et al., 2014; Yoshizawa et al., 2014). In contrast to normal culture conditions (no treatment; NT), GS consistently induced co-elution of both factors in high molecular weight fractions (around 1 MDa) of gel filtration chromatography analysis (Fig.3g). This observation suggests that both factors may be part of a multiprotein complex formed under these conditions. As further evidence of this link, SirT7 was able to immunoprecipitate endogenous mH2A1.1 specifically under GS (Fig.3h, lanes 3-4), an interaction that was abrogated by N189 mutation (Fig.3h, lanes 4-6).

### SirT7 binds to distal intergenic regions upon GS in a mH2A1-dependent manner

While the levels of total mH2A1 and mH2A1.2 decreased upon GS, we observed a consistently specific increase of mH2A1.1 under the same conditions, suggesting a direct role for this isoform in the GS response (Fig.4a). To understand the functional relationship between the two factors under GS we next studied the distribution of mH2A1 in Wt and SirT7-deficient primary mouse embryonic fibroblasts (MEFs) under NT and GS. While we observed a global enrichment of mH2A1 in Wt cells upon GS compared with NT conditions, this effect was abrogated in SirT7-deficient cells (Fig.4b). SirT7 ChIP-seq analysis showed that GS boosted SirT7 localization to chromatin (Fig.4c), a finding also supported by biochemical fractionation analysis of SirT7 (Supplementary Fig.3a). Interestingly, 72.4% (1711 of 2638) of SirT7-associated genes were also enriched by mH2A1. SirT7 accumulated mostly in intergenic regions (transcription start site [TSS] ± 50-500Kb) rather than in proximal promoters (Fig.4d). GREAT analysis associated these regions with specific genes (Fig.4e and Supplementary 3b), suggesting that the sequences correspond to gene regulatory regions. The average distance from these sites to the associated genes ranges between 50 Kb and 500 Kb upstream or downstream of the coding region. Many of the SirT7-associated genes mapped by GREAT were associated by KEGG analysis with cell signaling, the majority of which directly or indirectly participate in regulating metabolic homeostasis. We also found an unexpected overrepresentation among these genes of those involved in the neural system, particularly those associated with neurotransmitter activity, axon guidance and signal transmission (Fig.4e).

**Fig.4.**
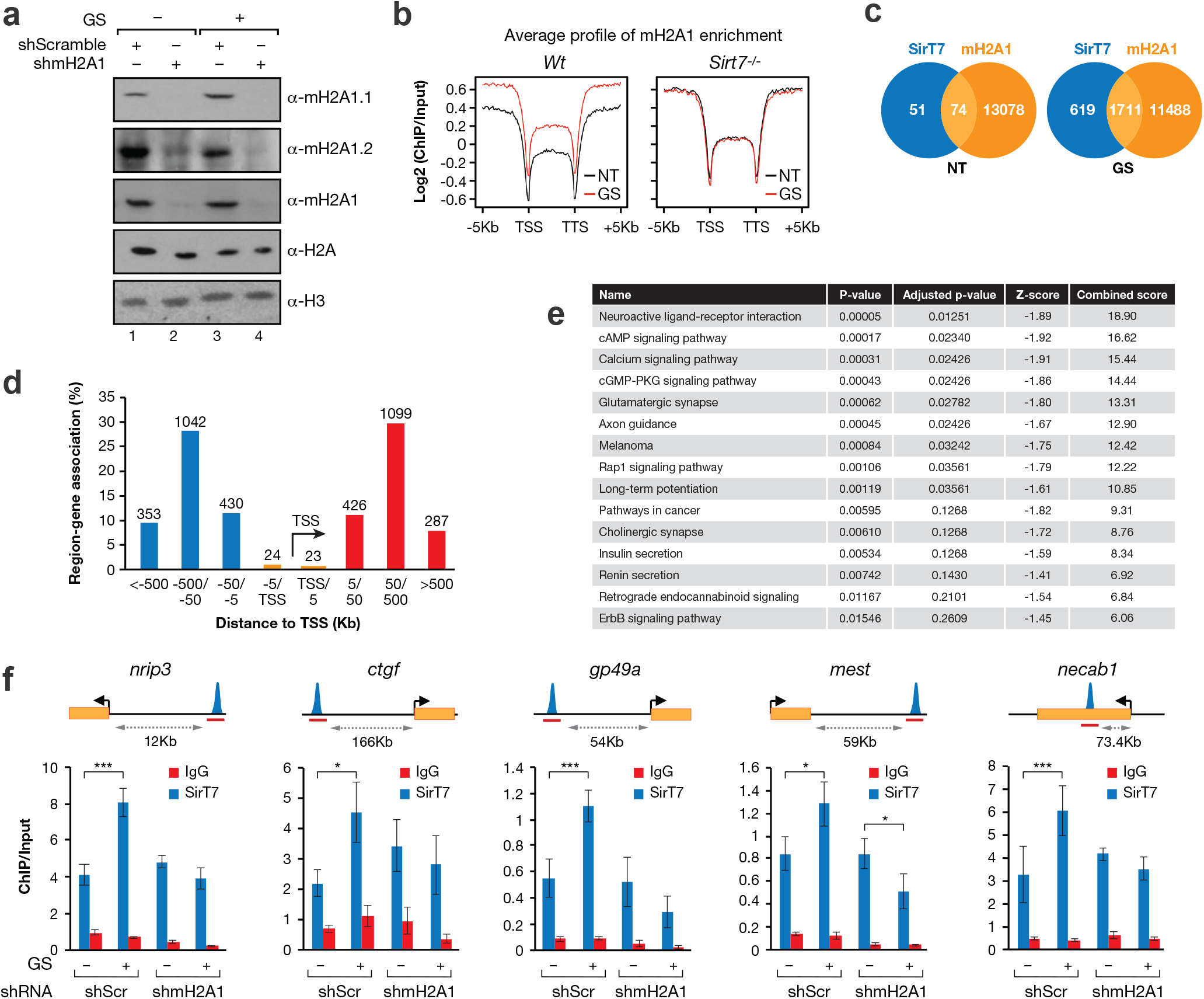
mH2A1 recruits SirT7 to distal intergenic regions associated to metabolic genes upon GS. **a**, Levels of three mH2A isoforms in whole-cell extract of NIH3T3 cells transfected with scramble shRNA or mH2A1 shRNA and cultured under normal or GS conditions. **b**, Plots showing average enrichment of mH2A1 at all genes in WT (left) or *SirT7*^-/-^ MEF (right) cells under no treatment (NT, black) or GS (red) conditions. Data expressed as the log_2_ ratio of RPKM normalized ChIP/INPUT signals. The gene body was scaled to 5 Kb for all genes, and the profiles extend 5 Kb upstream and 5 Kb downstream of the TSS and TTS (respectively) at a resolution of 100 bp. **c**, Venn diagrams showing the intersection of SIRT7-associated genes with mH2A1-enriched genes in WT cells under no treatment (NT, left) or GS (right) conditions. SIRT7-associated and mH2A1-enriched genes were derived from GREAT analysis (Supplementary Fig.3b and Methods). **d**, Distribution of sites occupied by SirT7 upon GS around the transcription start site (TSS). The values for each bin from the TSS are shown above each bar (TSS-5 Kb, 5-50 Kb, 50-500 Kb and >500 Kb). **e**, KEGG cell signaling pathways for SirT7-associated genes mapped by GREAT analysis under GS in MEF cells. The signaling pathways were ranked by their combined score (combination of Fisher’s exact test p-value and the z-score of the expected rank) provided by Enrichr analysis tool (Chen et al., 2013). **f**, SirT7 ChIP-qPCR analysis of SirT7 binding sites associated with mH2A1 enrichment at distal regions upon shRNA-mediated downregulation of mH2A1 under normal and GS conditions in NIH 3T3 cells. The amplified regions (red) and their distance to each gene are indicated in the upper part of the figure. Each SirT7 ChIP was normalized with respect to its own input, and error bars represent the standard error of the mean (SEM) from four independent experiments. Probabilities are those associated with a two-tailed t-test (*: p<0.05; **: p<0.01).

Confirming our previous results, the recruitment of SirT7 to these regions upon GS was mH2A1-dependent, since its shRNA-driven downregulation abrogated SirT7 enrichment (Fig.4f). These results indicate that, upon GS, mH2A1.1 recruits SirT7 to the intergenic regions, and suggest that SirT7 has a direct role in auto-mADPRTion.

### The SirT7/mH2A1 regulatory axis regulates the expression of a subset of genes upon GS

To clarify the mode of action of SirT7 in the genes regulated by SirT7/mH2A1 when they are stressed, we performed RNAseq in *Wt* and *Sirt7*^-/-^ MEFs under NT and GS as for the ChIP-seq analyses. Of the 1711 mH2A-containing regions occupied by SirT7, GS induced the upregulation and downregulation of 257 and 427 nearby genes, respectively, in *Wt* MEFs (Fig.5a-b). This up- or downregulation upon GS was SirT7-dependent in the case of 143 and 277 genes, respectively (Fig.5b). There was an increase in the levels of mH2A1 around these genes upon GS that was abrogated in SirT7-deficient cells (Fig.5c, Supplementary Fig.4). Among these genes we identified some that encoded growth factors, signaling transduction mediators involved in metabolism and differentiation, hormone receptors, enzymes, transcription factors, extracellular factors and cytokines. Many of these genes are directly or indirectly associated with the second messenger regulatory system, including the Ca^+2^-dependent pathways, and the cAMP/adenylate cyclase or cGMP-PKG signaling pathways. We also identified many factors associated with G-protein signaling, such as the G-protein-coupled receptor signaling pathways, some of which have been linked to regulation of cAMP signaling in neural systems. Genes downregulated upon GS were involved in cAMP signaling pathway factors, insulin secretion, proliferation and cell differentiation, while upregulated genes were mainly associated with cGMP-dependent protein kinase and Rap1 signaling, lipid metabolism and mobilization, and, to a much lesser extent, cAMP/adenylate cyclase signaling (Supplementary Fig.5). Among these up- or downregulated factors we identified, for instance, the growth factor CTGF, the adenylate cyclase regulator ADRA2A, the GTPase GBP6, the histone deacetylase HDAC9, the calcium-binding signaling protein NECAB1, and the transcription factors FOXA2 and MEF2C. CTGF was particularly interesting, given its key role in differentiation (Guney et al., 2011), proliferation (Riley et al., 2015), glucose homeostasis (Dai et al., 2016; Lam et al., 2003; Liu et al., 2007), IGF-1 (Zhou et al., 2008) and TGF-β signaling (Abreu et al., 2002; Tsai et al., 2018), and its link to calorie restriction (CR) (Forrester et al., 2014) and aging (Ungvari et al., 2017).

**Fig.5.**
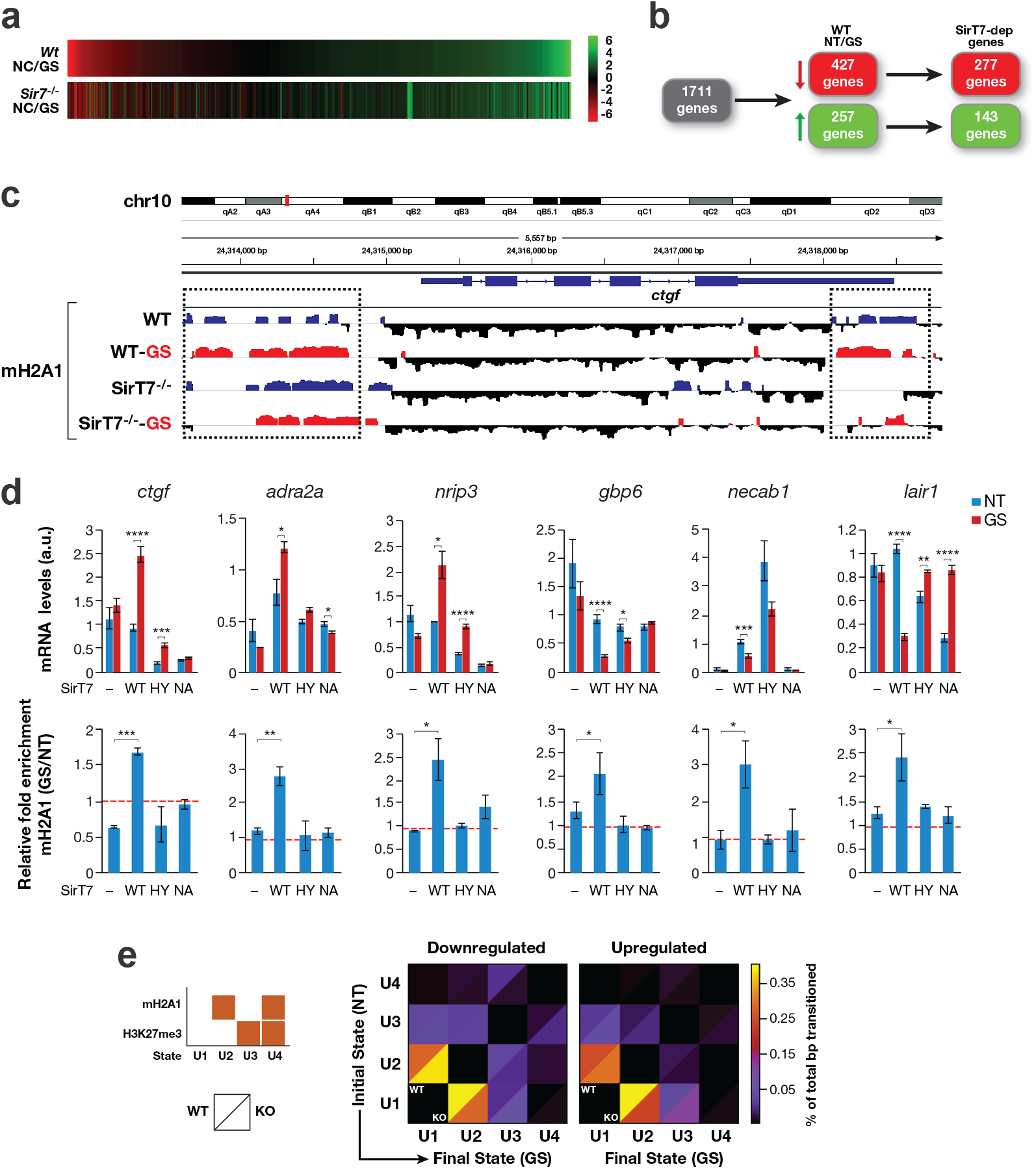
SirT7 dual activity promotes mH2A1 enrichment in the promoters of associated genes resulting in transcription upregulation or downregulation. **a**, Heatmap showing RNA-expression changes relative to no treatment (NT, as log_2_ magnitude of difference between GS and NT) conditions in WT and SirT7-deficient MEFs. **b**, Schematic diagram showing the bioinformatic pipeline applied to RNAseq data to filter genes associated with SirT7/mH2A1 in *Wt* and *Sirt7*^-/-^ MEF cells treated under GS or not (NT). Analysis was restricted to genes that: 1) were associated with SirT7 via GREAT and were mH2A1-enriched; 2) showed a log_2_ FC of expression between WT-NT and WT-GS of > 0.6; and 3) a difference between WT and KO log_2_ FC (NT *vs*. GS) of > 0.3 (Table S3). **c**, Integrative Genome Viewer (IGV) of mH2A1 ChIP-seq signals across *ctgf*, a gene differentially enriched in mH2A1 upon GS in *Wt* MEFs (red, positive values) compared with NT (blue, positive values), and whose expression is controlled by SirT7 under the same stress conditions. **d**, Upper panel: RT-qPCR analysis of genes regulated by SirT7 upon GS and NT. The expression of SirT7 in *SirT7*^-/-^ MEFs was rescued by retroviral-mediated gene transfer of SirT7 WT, H187Y (HY), N189A (NA), and empty vector (-). Error bars represent the standard error of the mean (SEM) from four independent experiments. Probabilities are those associated with two-tailed t-tests (*: p<0.05; **: p<0.01; ***: p<0.005; ****: p<0.001). Bottom panel: relative mH2A1 enrichment (GS *vs*. NT) by ChIP-qPCR analysis at specific regions around the TSS of the indicated genes (*gbp6*: −6 Kb; *necab1*: −3 Kb; *lair1*: −27.5 Kb; *ctgf*; +600 bp; *adra2a*: −5 Kb; *nrip3*: +7 Kb). Error bars represent SEM from three independent experiments. Probabilities are those associated with one-way ANOVAs (*: p<0.05; **: p<0.01; ***: p<0.005). **e**, Heatmaps showing chromatin state transitions induced by GS. Chromatin state transitions induced by GS in WT and SirT7-deficient cells were collected and normalized with respect to the total number of base pairs that transitioned to another state (as the percentage of all transitioned base pairs). Bottom left: Depiction of the relative location of WT and KO data in each heatmap square. Bottom right: State map illustrating the specific combination of mH2A1 and/or H3K27me3 in the four chromatin states defined in the analysis. U1: without H3K27me3 or mH2A1; U2: enriched by mH2A1; U3: enriched by H3K27me3; U4: enriched by H3K27me3 and mH2A1.

Reconstitution experiments with SirT7 WT, H187Y and N189A in Sirt7^-/-^ MEFs revealed that both enzymatic activities of SirT7 are required to produce the GS-associated expression profile of the tested genes, as well as the associated enrichment of mH2A detected in these genes upon GS (Fig.5d). Consistent with a direct role of SirT7 in these genes through binding to the distal regulatory regions, we also observed SirT7-dependent H3K36 deacetylation in these distal sequences, which was dependent on both catalytic activities (Supplementary Fig.6). These lines of evidence suggest a link between this deacetylation event and mH2A gene enrichment. As mH2A1 was previously associated with H3K27me3 (Buschbeck et al., 2009), we also analyzed the correlation between this mark and mH2A/SirT7 through a global unbiased analysis of chromatin state changes from NT to GS conditions in *Wt* and *Sirt7*^-/-^ MEFs. We defined four states (U1-U4), depending on the appearance or disappearance of mH2A and/or H3K27me3 in these sequences upon GS. We observed a positive effect of SirT7 on mH2A recruitment upon GS in upregulated and downregulated genes when H3K27me3 was not present. In contrast, when mH2A1 and H3K27me3 were both present, SirT7 had the opposite effect on mH2A enrichment (Fig.5e). These results suggest an antagonistic interplay between SirT7 and H3K27me3 in determining mH2A enrichment upon metabolic stress (Fig.5e).

### SirT7 dependence on the expression of these genes is recapitulated in *in vivo* mouse models of CR and aging

To confirm *in vivo* the contribution of SirT7 to GS response, we generated a CR model in *Wt* and *Sirt7*^-/-^ mice (Fig.6a). We studied the expression of the SirT7/mH2A-associated genes we had identified earlier (Fig.4–5) in the livers of these mice. The experiments fully replicated our results in cell culture with GS, providing further evidence of a key role for the SirT7/mH2A axis in glucose/CR signaling (Fig.6b).

**Fig.6.**
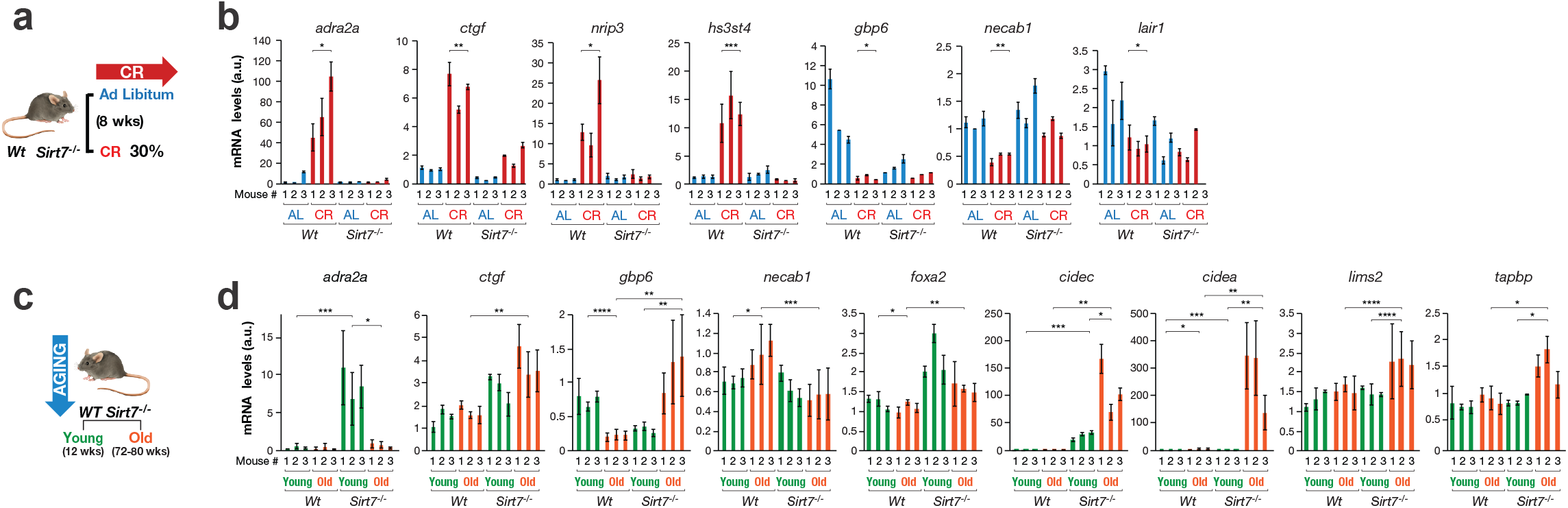
SirT7 and mH2A1 axis plays a key role in calorie restriction and aging. **a**, Model validation studies in WT and SirT7 KO mice fed *ad libitum* or calorie-restricted (CR, 30%) for 8 weeks. **b**, RT-qPCRs analysis of the indicated genes in liver samples from *Wt* and *Sirt7*^-/-^ mice fed *ad libitum* or CR. Three animals were analyzed for each condition. Each quantification was generated from three replicates. Probabilities are those associated with one-way ANOVAs (*: p<0.05; **: p<0.01; ***: p<0.005). **c**, Aging studies carried out in liver samples taken from young (12 weeks) and old (72-80 weeks) *Wt* and *Sirt7*^-/-^ mice. **d**, Gene expression measured by RT-qPCR in liver samples from young and old *Wt* and *Sirt7*^-/-^ mice. Three animals were analyzed for each condition. Each quantification was generated from three replicates, and probabilities are those corresponding to t-tests. (*: p<0.05; **: p<0.01; ***: p<0.005).

Given the established link between CR and aging and the aging-accelerated phenotype observed in *Sirt7*^-/-^ mice (Vazquez et al., 2016), we next tested whether SirT7 is responsible for the aging-associated expression profile of these genes. To this end, we analyzed their expression levels in the livers of young (12 weeks) and old (72-80 weeks) *Wt* and *Sirt7*^-/-^ mice (Fig.6c). Strikingly, the gene-specific expression changes observed upon aging were dependent on SirT7 in the vast majority of the genes tested (Fig.6d). We also observed a similar dependence on expression of key metabolic regulators linked to aging and CR, such as *cidec* and *cidea* (Abreu-Vieira et al., 2015; Baur et al., 2006; Magnusson et al., 2008). The aging-associated expression of some mouse aging markers (Maegawa et al., 2017) was also dependent on SirT7. Overall, our findings not only suggest a key role for SirT7 in aging through gene expression regulation of second metabolism signaling, but also that this role is directly related to calorie restriction (Fig.7).

**Fig.7.**
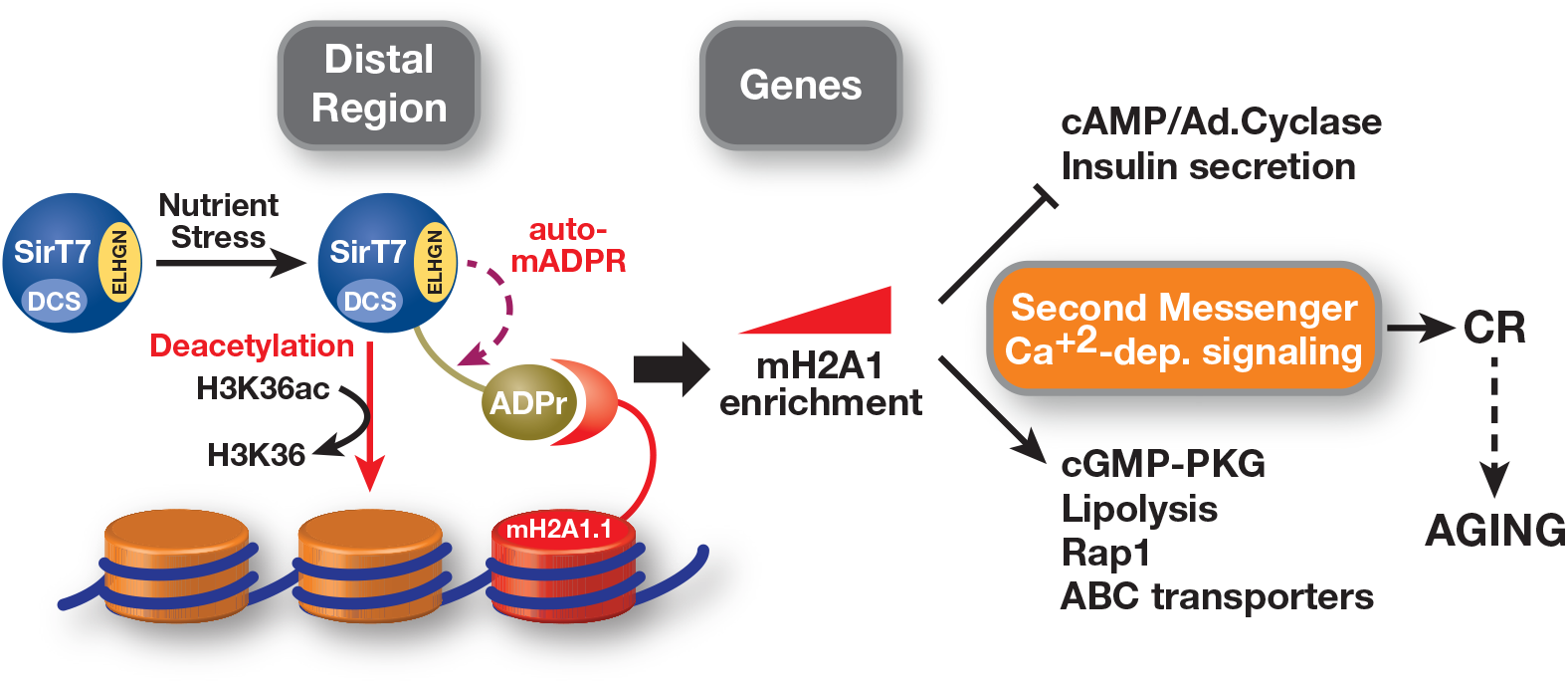
SirT7/mH2A1 regulatory axis links glucose starvation with calorie restriction and aging. Model proposed for the dual SirT7/mH2A regulatory axis in GS, CR and aging. Nutrient stress induces SirT7 auto-mADPRT, which leads to mH2A-dependent recruitment of SirT7 to distal regulatory regions and subsequent mH2A enrichment around the associated genes. This axis plays a key role *in vivo* in CR and aging by modulating key signaling pathways.

## Discussion

We report for the first time ADPRT activity in SirT7 and identify a novel ADP-ribosylation domain containing the ELGHN motif. The direct link between this modification, SirT7 dynamic distribution, and mH2A1 binding in response to GS provides a functional explanation of sirtuin enzymatic duality and suggests a key role for SirT7 in CR signaling and aging (Fig.7).

The catalytic duality of sirtuins has been widely debated since it was hypothesized that all sirtuins may have the ability to transfer ADP-ribosyl moieties to target proteins (Frye, 1999; Tanny et al., 1999). Originally described as a potential enzymatic alternative to the main deacylation/deacetylation activity, it was assumed to act alongside the primary activity at the same catalytic site. This perception arose primarily because the ADPRT activity of some of the sirtuins tested was considerably weaker and less specific than the deacetylase activity, and was dependent on the presence of an acetylated peptide (Imai et al., 2000; Tanny et al., 1999). Moreover, the dependency of this activity on the deacetylation reaction of yeast Sir2 was also reported, suggesting that it may be a side effect of primary catalytic deacylation or a technical artifact linked to the ability of sirtuins to hydrolyze NAD^+^ as an intermediate step before deacetylation (Moazed, 2001) It is of particular note that the vast majority of this evidence was based on *in vitro* ADP-ribosylation assays and was not validated by other techniques. In this sense, we report the first MS analysis of sirtuin ADPRTion and describe a consistent and reproducible pattern of N189-dependent ADPRTion in class IV sirtuins, the common lineage of SirT6 and SirT7 from early eukaryotes to mammals. We not only demonstrate that the role of the ELGHN motif in this activity is a highly conserved one, but also that the catalytic site is physically separated, being located in a different structural domain. Our studies suggest that N189 could play an important role as the first acceptor of the ADP-ribose molecule generated by E185 before its transfer to the final target. This is supported by the direct interaction with E185 and the different behavior of N189Q compared with WT, despite its close structural similarity (Supplementary Fig.1e). Given the wide distribution of these modified residues across the surface of SirT7, the auto-modification probably involves the intra-modification of SirT7 oligomeric units within the SirT7 complex, similar to what has been described for PARP1 (Altmeyer et al., 2009).

Although auto-ADPRTion had already been described in the early studies of sirtuin, its role has never been properly characterized. Our report suggests a direct role of this modification in SirT7 global distribution as well as in its chromatin-binding dynamics. PARP1 auto-ADPRTion was also proposed to regulate PARP1 chromatin binding dynamics (Muthurajan et al., 2014). As in the case of SirT7, mH2A1.1 has been also involved in PARP1 recruitment, although in this case it was linked to PARP1 inactivation (Ouararhni et al., 2006; Sun and Bernstein, 2019). Although our work has focused on auto-ADPRTion, the high degree of conservation within the class IV sirtuins and the considerable magnitude of the cavity, which makes it able to accommodate a wide range of proteins (Fig.1e, Supplementary Fig.1d), suggest that this novel domain may target other substrates.

In contrast to what was reported for SirT6-dependent ADPRTion of PARP1 and KAP1 (Mao et al., 2011; Van Meter et al., 2014), SirT6 and SirT7 auto-mADPRTion do not depend on any key residues involved in the primary deacetylation domain, such as SirT6-S56/SirT7-S111 or SirT7-S115. This suggests that there may be two types of sirtuin-related ADP-ribosylation events, one that takes place at the primary catalytic site and that is probably directly associated with the main deacetylation reaction, and another that is dependent on the ELGHN motif including auto-ADP-ribosylation and modification of other protein targets. This hypothesis suggests an interesting line of future work to confirm its validity, and thereafter to define the extent and the specific range of substrates associated with each of these ADP-ribosylation events.

Another major finding of our work is the first evidence of a direct functional relationship between sirtuins and mH2A through this ADPR activity. This link is supported by the ability of mH2A1.1 to bind to the sirtuin product O-acetyl-ADP ribose (Pazienza et al., 2014). Our results demonstrate that GS triggers SirT7 relocalization mainly to distal regulatory regions mediated by mH2A1.1. These regions do not appear to be canonical enhancer regions because they are not enriched in enhancer-specific histone marks (data not shown). In turn, SirT7 promotes increased mH2A1 occupancy levels in the nearby gene, which results in the transcriptional activation or repression of the gene (Fig.5). The fact that SirT7 seems to induce both up- and downregulation in these genes suggests that its main role in this functional context is to promote mH2A enrichment, which in turn may regulate positively or negatively gene expression. The analysis of chromatin states suggests that H3K27me3 is antagonistic to the interplay between SirT7 and mH2A. This intriguing observation deserves further study. The mechanism by which SirT7 promotes this long-range effect on mH2A enrichment around genes is unclear, and it may involve chromatin looping, as occurs in enhancer regions (Calo and Wysocka, 2013). This possible mechanism is consistent with the recently established link between SirT7 and the nuclear lamina and chromatin organization (Vazquez et al., 2019). Our results suggest that the role of SirT7 in these genes requires both enzymatic activities, which further highlights the essential role of sirtuin duality in this regulatory mechanism. In agreement with this, we detected active deacetylation of H3K36 in the distal regulatory regions upon the arrival of SirT7, which may favor the mechanism of mH2A enrichment.

This study also suggests that the Sirt7/mH2A axis regulates a wide range of factors under GS. We were able to recapitulate the GS stress conditions found in primary MEFs from livers of calorie-restricted mice. Consistent with previous studies, we detected gene expression profile differences associated with caloric restriction (e.g., ADRA2A, CTGF and LAIR1) (Ogawa et al., 2011; Swindell, 2009; Yamamoto et al., 2009), but also new potential markers involved in obesity (NRIP3) (Wen et al., 2019) and immune response (HS3ST4, GBP6) (Kommadath et al., 2017; Stacey et al., 2017). Remarkably, their stress-dependent expression response profile was SirT7-dependent, confirming the function of SirT7 as a nutrient sensor in response to calorie-restricted diet. The observed overrepresentation of second messenger signaling among these factors, particularly that related to calcium-associated signaling, is quite surprising because it suggests this is a major target of the SirT7/mH2A signaling axis. Considering the wide range of general and tissue-specific functions regulated by these pathways, the functional relevance of this regulatory axis may extend beyond glucose metabolism. cAMP/adenylate cyclase signaling was shown to regulate SirT6 activity, which suggests regulatory feedback may operate in this context (Kim and Juhnn, 2015). Another significant observation of our work is that SirT7 is important in the aging-associated changes linked to glucose homeostasis regulation. Although SirT7 was previously associated with aging at different levels, from genome stability and structure to ribosomal biogenesis and mitochondrial function (Ford et al., 2006; Ryu et al., 2014; Vazquez et al., 2016; 2019), our study is the first to establish a link between SirT7 and CR signaling and to demonstrate a direct effect of SirT7 on the expression profile of this signaling during aging. This link is supported by the *in vivo* metabolic alterations detected in SirT7-knockout mice, such as reduced IGF-1 plasma levels and hepatic steatosis (Ryu et al., 2014; Vazquez et al., 2016), and the significant upregulation of insulin signaling and lipid metabolism pathways, Cidec and Cidea (Yoshizawa et al., 2014), as well as other aging markers (Fig.6d) (Maegawa et al., 2017). Interestingly, many genomic structural alterations caused by aging in mouse livers seem to take place in regions 50-500 Kb upstream and downstream of the TSS of genes, which resembles SirT7 binding profile upon GS. Some of the factors regulated by SirT7, such as CTGF or FoxA2, have been directly implicated in calorie-restriction signaling and aging (Estep et al., 2009; Panowski et al., 2007; Ungvari et al., 2017; Whitton et al., 2018). Further work is required to assess the contribution of these targets to the accelerated aging phenotype associated with SirT7.

Our work not only offers a new insight into sirtuin duality and highlights the crucial role of sirtuins in the epigenetic regulation of metabolism and calorie-restriction signaling, but also provides a broader perspective on the dynamics and regulation of sirtuin function.

## ACKNOWLEDGMENTS

We are grateful to Dr. H.F. Willard, Dr. E. Verdin, and Dr. J. Gurdon for sharing reagents, and to Dr. L. Pardo for access to computing facilities. We also thank the members of the Vaquero laboratory for stimulating discussions. This work was supported by Spanish Ministry of Economy and Competitiveness - MINECO (SAF2011-25860, SAF2014-55964R, SAF2017-88975R to A.V.; SAF2016-77830R to M.O.) cofunded by FEDER funds/European Regional Development Fund (ERDF) - A Way to Build Europe, the Catalan government agency AGAUR (2014-SGR400, 2017-SGR148 to A.V.), a grant from Rutgers Human Genetics Institute of New Jersey (J.T., L.S.), the German Center for Cardiovascular Research (DZHK) and the Deutsche Forschungsgemeinschaft (DFG SFB TRR81 A02 and SFB 1213 TP B02 to A.I., T.B.). We also thank the CERCA Programme/Generalitat de Catalunya for institutional support. Proteomic analyses were performed in the IDIBELL and CRG Proteomic Units, both of which are part of Proteored PRB3 and are supported by grant PT17/0019 from the PE I+D+i 2013-2016, funded by ISCIII and ERDF.

## AUTHOR CONTRIBUTIONS

N.G.S and A.V. conceived the study and designed the experiments. N.G.S performed most of the experiments. A.V. supervised the experiments, coordinated the work and wrote the paper. J.K.T, B.N.V., J.T., and L.S. generated the SIRT7 KO MEFS, performed and analyzed the ChIP-seq and RNA-seq experiments, and the aging mouse model. M.E.-A. provided support for the ChIP and qPCR experiments. M.O. performed all the structural analyses. A.I. and T.B. carried out the calorie-restriction procedure, and provided KO MEFs and reagents. J.M.-S., E.S. and C.D.L.T. did the MS/MS analyses. M.B. and S.H. provided reagents and expertise. M.E. participated in the design, generation and analysis of the RNA-seq experiments. All authors participated in the discussions leading to the production of this paper.

