## Supplementary material for "SirT7 auto-ADP-ribosylation regulates glucose starvation response through macroH2A1.1": Methods

### SirT7 Computational Molecular Model

For *in silico* SirT7 structural model, an initial homology model was constructed for human SirT7 using the coordinates of the determined X-ray crystal structure of human SirT6 (PDB ID: 5M6F, 1.87 Å of resolution, 42% of sequence identity)[38]. Modeller 9.12 was used to model the non-determined regions of the protein[39]. The side chain conformations for non-conserved residues were positioned according to Scwrl 4 (Krivov). The protein was embedded in a tip3p water box. The initial system was energy minimized, subjected to 10 ns of molecular dynamics equilibration and finally to a production stage extending to 150 ns. All the simulations were performed with GROMACS 5.0 simulation package[40]. The molecular model of SirT7 was used as the initial model to introduce H187Y and N189A mutations. For comparison purposes, SirT1-5[41-45] structures were superimposed to SirT6 structure and SirT7 structural model using PyMOL (The PYMOL Molecular Graphics System, Version 1.8 Schrödinger, LLC.).

### *In vitro* enzymatic assays

ADP-ribosylation assays were performed in a total volume of 50 µl in 50 mM Tris-HCl, pH 7.8, 150 mM NaCl, 10 mM DTT, 1 µM unlabeled NAD+, 8 µCi of [$^{32}$P]-NAD$^+$ (800 Ci/mmol; PerkinElmer) incubated with 3-5 µg of recombinant protein from *E. coli* (BL21) or purified from HEK293F cells. The reactions were incubated at least for 2 hours at 37ºC. Reactions were stopped by adding 5X Laemmli sample buffer (supplemented with 10% 2-Mercaptoethanol) to a final concentration of 1X and then separated by SDS-PAGE. Following gel electrophoresis proteins were then transferred from gel to PVDF membrane (Millipore) in transfer buffer (500 mM glycine, 50 mM Tris-HCl, 0.01% SDS, 20% methanol) at 400 mA for 1 hour on ice. PVDF membranes were then stained with Coomassie Brilliant Blue (Sigma-aldrich) and the incorporation of ADP-ribosylation into proteins was visualized by autoradiography.

Deacetylation assays were performed in HDAC Buffer (10 mM Tris-HCl pH 7.8, 150 mM NaCl, 1mM DTT, 10% Glycerol) with 5 mM of $NAD^+$ (Sigma-Aldrich) in the presence of 1-3 µg of SirT7 and ~0.5 µg of hyperacetylated histone substrates for 3 hours at 37ºC. Hyperacetylated histone substrates were acid-extracted from HeLa cells treated with 5 µM TSA and 5 mM nicotinamide. The reactions were stopped by adding 5x Laemmli sample buffer (supplemented with 10% 2-Mercaptoethanol) to a final concentration of 1X of Laemmli sample buffer and then separated by SDS-PAGE. The acetylation status was determined by Western-blot analysis using specific antibodies.

**Recombinant protein purification**

Human cDNAs of SirT6 or SirT7 were cloned into the pET-30b expression vector (Novagen, Germany), containing a hexahistidine tag at the C terminus. The constructs were expressed in *E.coli* BL21 (DE3) grown at 37ºC unit $OD_{600}$=0.5 and then induced with 0.4 mM IPTG for 4 hours. Harvested cells were resuspended in lysis buffer (20mM Tris pH8.0, 100mM NaCl, 10mM EDTA pH8, 0.5% NP40 and 2% sarkosyl) and lysed by sonication. After centrifugation at 15,000 x *g* in a Sorvall SS-34 rotor, the cleared lysated were loaded onto Ni-NTA agarose resin (Qiagen) by gravity flow. After several washes with lysis buffer and BC-500 (20 mM Tris, pH 8.0, 500 mM KCl, 10% glycerol, 1 mM EDTA, 1 mM DTT, 0.1% Nonidet P-40, 0.2 mM PMSF), rSirT7 was eluted using 100 mM Imidazol resuspended in BC500. Recombinant proteins were then kept at -80ºC.

Preparation of recombinant histones were performed from *E.coli* Bl21 (DE3) cells as described elsewhere[46]. Briefly, *E.coli* Bl21 (DE3) cells were transformed with pET-histone expression plasmids of the *Xenopus laevis* histone genes for H2A, H2B, H3, H4 and grown at 37°C to a density of $OD_{600}$=0.5-0.6 and then induced with 1 mM IPTG for 4 hours. The bacteria pellet was washed 3 times with TW buffer (50 mM Tris-HCl, pH 7.5, 100 mM NaCl, 1 mM EDTA, 1 mM benzamidine, 5 mM 2-mercaptoethanol, 1% v/v Triton X-100) and then resuspended in unfolding buffer (7M guanidinium HCl, 20 mM Tris-HCl, pH 7.5, 10 mM DTT) for 1 hour at room temperature. Cell debris was removed by centrifugation at 20 ºC and 23,000 x *g*, and the supernatant containing the unfolded proteins was loaded into a Gel filtration column XK-50 (Pharmacia,

Uppsala, Sweden), packed with Sephacryl S-200 high-resolution gel filtration resin (Pharmacia). Elutions profiles were recorded at a wavelength of 280 nm, collected and visualized by 18% SDS-PAGE. Proteins were dialyzed (dialysis bags with a cutoff of 6-8 kDa) in 7M urea, 20 mM sodium acetate, pH 5.2, 2M NaCl, 5 mM 2-mercaptoethanol, 1 mM EDTA and then loaded into TSK SP-5PW HPLC column. For the refolding of histone octamers, unfolded histone proteins were dialyzed at 4ºC against at least three changes of 2L of refolding buffer (2M NaCl, 10 mM Tris-HCl, pH 7.5, 1 mM EDTA, 5 mM 2-mercaptoethanol). The purity and stoichiometry of the fractions were checked on an 18% SDS-PAGE. Histone octamers were concentrated to 3-15 mg/mL, adjusted to 50% (v/v) glycerol, and stored at -80ºC.

**Statistical analysis**

The represented values show means of at least three of more independent experiments (n≥3 with error bars representing standard error of mean (SEM) unless otherwise specified. Data were analysed using two-tailed student´s *t*-test (ADP-ribosylation and deacetylation assays, RT-qPCRs from rescue experiments in MEFs), and for multiple comparisons the *P* value was calculated using Microsoft Excel software by two-way ANOVA (RT-qPCR in aged mice and from mice under caloric restriction and *ad libitum* feeding).

**Cells transfections and treatments**

MEFs derived from WT and SirT7 knockout mice were generated from day 13.5 embryos by standard methods[47]. For retroviral infection of primary MEFs and NIH3T3 cells, Platinum-A cells were transiently transfected using polyethylenimine (PEI). After 6 hours of transfection, the media was replaced with fresh media for 48 hours. The collected viral suspension (culture media) was filtered using a syringe filter containing 0.45 µm in diameter pore filter. Cells were incubated with virus-containing supernatant in the presence of 8 µg/ml polybrene. In the case of shRNA in NIH3T3 cells, after 48 hours of infection, infected cells were selected for 72 h with puromycin (2.0 µg/ml). Plasmids encoding mouse shRNAs of macroH2A isoforms were obtained from Addgene (Cambrige, MA, USA): pSUPER retro puro macroH2A1 shRNA (#30517), pSUPER retro puro macroH2A2 shRNA (#30518), pSUPER retro puro Scr shRNA (#30520). HEK293F cells were transiently transfected with plasmids using 3 uL of polyethylenimine (PEI) 1mg/mL per µg of DNA. The following plasmids were used for transfection: pcDEF-Flag-SirT7 (wild-type, H187Y, N189A and N189Q), pCMV6-SirT7-GFP (Origene RG205658).

For glucose starvation, cells were treated 24 hours prior harvesting with glucose-free Dulbecco's modified Eagle's medium (DMEM) (GIBCO) containing 10% fetal bovine serum. For oxidative stress, cells were treated with 1mM $H_2O_2$ for 1h before harvesting. Cells were irradiated by exposure in the following conditions: a) UV irradiation at 254nm (80 J/m$^2$) in a Stratagene Stratalinker 2400 UV (Agilent technologies); b) Gamma irradiation delivered using an aluminium filter at 100kVp in a Faxitron Cabinet X-ray system (Faxitron corp.).

**Western blot and immunoprecipitations**

For western blot and co-immunoprecipitations, whole-cell preoteins extracts were prepared according to the Dignam protocol[48]. Protein immunoprecipitations were performed with the respective antibodies conjugated to agarose beads (Millipore) for 6 hours with gently agitation at 4 ºC. The affinity-purified protein complexes were gently washed at least 3 time with BC300 (20 mM Tris, pH 8.0, 300 mM KCl, 10% glycerol, 1 mM EDTA, 1 mM DTT, 0.1% Nonidet P-40) and eluted by either with 3XFLAG peptide (Anaspec) or by acidification using a buffer containing 0.2 M glycine, pH 2.3

Co-immunoprecipitations were performed using FLAG-agarose (Sigma), HA-agarose (Sigma), Myc antibody (Cell Signaling), and Protein G Agarose (Millipore). Densitometric analysis of the western blots was performed with Quantity One software (Biorad). The following antibodies were used for western blotting: anti-Flag M2 Antibody (Sigma-Aldrich; F1804); anti-Fibrillarin (Santa cruz, B-1, Sc-166001); anti-H3 (Cell Signaling, #9715); anti-tubulin (Abcam, ab-6160); anti-SirT7 (Cell Signaling, D3K5A); anti-macroH2A1.1 (Cell Signaling, D5F6N); anti-macroH2A1.2 (Cell Signaling, #4827); anti-macroH2A1 (abcam, ab37264); anti-macroH2A.2 was gifts of Marcus Buschbeck; anti-histone H2A

(Cell Signaling, #12349), anti-HA (Sigma-Aldrich, H6908); anti-myc (Cell Signaling, #2276), anti-H3K18ac (Cell Signaling, #9675).

**Chromatin fractionation**

To determine the amount of SirT7 proteins in cytoplasm, nucleoplasm and tight chromatin fraction, NIH3T3 cells were transfected with SirT7-HA WT, H187Y, N189Q and N189A. cytoplasmic and nucleoplasm fraction was obtained with the Dignam protocol extraction (fractions A and C). The chromatin pellet was washed with BC-500 and then solubilized in Laemmli buffer and sonicated with Branson 250 sonicator.

**Mass spectrometric analysis**

**Protein digestion analysis**

Protein extracts were precipited with cold acetone (o/n, -20ºC) and the protein pellets were ressuspended in 20 µL of 200 mM ammonium bicarbonate (ABC) + 8M urea solution. Samples were reduced with DTT (10 mM, 30 ºC, 60 min), and alkylated with iodoacetamide (20 mM, 25 ºC, 30 min). The samples were diluted with $NH_4HCO_3$ to reduce urea concentration below 1M, and digested with trypsin (1:10 enzyme:protein, 32ºC, 16h, Promega). After digestion, the peptide mix was acidified with formic acid and then desalted with a MicroSpin C18 column (The Nest Group) prior to LC-MS/MS analysis.

**Mass spectrometric analysis**

The peptide samples were analyzed using a Lumos Orbitrap mass spectrometer (Thermo Fisher Scientific, San Jose, CA, USA) coupled to an EASY-nLC 1000 (Thermo Fisher Scientific (Proxeon), Odense, Denmark). Afterwards peptide were loaded directly onto the analytical column and were separated by reversed-phase chromatography using a 50-cm column with an inner diameter of 75 µm, packed with 2 µm C18 particles spectrometer (Thermo

Scientific, San Jose, CA, USA) and a binary solvent system of 0.1% formic acid in H2O (Solvent A) and 0.1% formic acid in acetonitrile (Solvent B). Chromatographic gradients started at 7% B with a flow rate of 300 nl/min and gradually increased to 22% B in 52 min and then to 32% B in 8 min and 95% B in 10 min. After each analysis, the column was washed for 10 min with 95% B.

The mass spectrometer was operated in positive ionization mode with nanospray voltage set at 2.4 kV and source temperature at 275°C. Ultramark 1621 for the was used for external calibration of the FT mass analyzer prior the analyses, and an internal calibration was performed using the background polysiloxane ion signal at m/z 445.1200.
The acquisition was performed in data-dependent acquisition (DDA) mode and full MS scans with 1 micro scans at resolution of 120.000 were used over a mass range of m/z 300-1500 with detection in the Orbitrap. Precursor automated gain control (AGC) was set to 2e5. Charge-state screening was enable, and precursors with +2 to +5 charge states and intensities >5e4were selected for tandem mass spectrometry (MS/MS). MS/MS acquisition method used was previously described[49]. A low-resolution and high-energy data-dependent HCD (acquired in the orbitrap with collision energy of 38% and a resolution of 3e4. AGC was set to 5e4, isolation window of 2 m/z and maximum injection time of 60 ms was used) was followed by high-quality HCD (acquired in the orbitrap with collision energy of 35% and a resolution of 12e4. AGC was set to 5e5, isolation window of 2 m/z and maximum injection time of 240 ms was used) and EThCD MS/MS when more than two ADP-ribose fragment peak (136.0623, 250.0940, 348.07091 and 428.0372) were observed in the low resolution HCD scan. For the EThCD fragmentation, the “use-calibrated charge-dependent parameter” option was selected, it was acquired in the orbitrap with a SA of 25% and a resolution of 12e4. AGC was set to 5e5, isolation window of 2 m/z and maximum injection time of 240 ms was used. Precursors masses previously selected for MS/MS measurement were excluded further selection for 60s and the exclusion window was set at 10 ppm. All data were acquired with Xcalibur software v4.1.31.9.

### Data Analysis

The data analysis was performed using two different software (Proteome Discoverer and Peaks). The Proteome Discoverer software suite (v2.2, Thermo Fisher Scientific) and the Mascot search engine (v2.5 Matrix Science[50]). The data were searches against a Swiss-Prot human database (as in April 2018, 20797 entries) plus a list of common contaminants [refined list from contaminants.fasta of MaxQuant] and all the corresponding decoy entries. All MS/MS spectra were deconvoluted by use of the MS Spectrum Processor. For peptide identification, a precursor ion mass tolerance of 10 ppm was used for MS1 level and the fragment ion mass tolerance was set to 0.05 Da for MS2 spectra. Enzyme specificity was set to trypsin, and up to five missed cleavages allowed. Oxidation of methionine, N-terminal protein acetylation and ADP-ribosylation were used as variable modifications whereas carbamidomethylation on cysteines was set as a fixed modification. The ADP-ribose modification was set differently for HCD and EThcD, as previously described[51]. This modification was set in the following residues: DEKNQRSTY. False discovery rate (FDR) in peptide identification was set to a maximum of 5%. The spectra analyzed using the PEAKS7 software[52] which as a default performs de novo peptide sequencing prior to database searches, in order to improve the accuracy of the results. Data were searched against the same reference Uniprot Homo sapiens database that was used in Proteome Discovered analysis. Trypsin (specific, up to four missed cleavages allowed and one non-specific cleavage) was selected for database searches, and no enzyme was chosen in de novo searches (up to 5 candidates per spectrum reported). The maximal mass error was set to 10 ppm for precursor ions and 0.05 Da for product ions. Carbamidomethylation was selected as a fixed modification, and methionine oxidation and ADP-ribose (+541.0611 Da at KESNCRDTQY) were set as variable modifications. The maximal number of modifications per peptide was set as five. The false discovery rate was set to 0.01 for peptides.

### Animal studies, calorie restriction and studies

For the Calorie restriction procedure, 8 weeks old male WT and SirT7$^{-/-}$ animals[53] were housed in individual cages and randomly allocated into *Ad Libitum* or calorie restriction (CR) group. The average of food eaten *Ad libitum* for each mouse was determined by weighing the remaining food on a daily basis for 1 week. The CR regimen was applied by feeding daily the animals with 70% of the calculated amount for 8 weeks. All animal experiments were performed in accordance with the Guide for the Care and Use of Laboratory Animals published by the National Institutes of Health[54] and were approved by the local authorities (RP Darmstadt; Germany).

For aging studies, livers of young (3 months) or old (18 to 20 months with an accelerated aging phenotype[28]) *Wt* and *Sirt7*$^{-/-}$ mice (male and female) were collected.

**RNA isolation, cDNA synthesis and RT-qPCRs.**

Total RNA was purified from mouse embryonic fibroblasts (MEFs) and liver using TRIzol reagent (Invitrogen, Carlsbad, CA, USA). The integrity and quality of RNA was evaluated on 1.2% agarose gel electrophoresis and quantified spectrophotometrically at A260nm. The cDNA was synthesized from 5 µg of total RNA with transcriptor first strand cDNA synthesis kit (Roche) according to the manufacturer's instructions. Real-time quantitative PCR was performed using the QuantStudio™ 5 Real-Time PCR System (ThermoFisher Scientific) with Sybr green PCR Master Mix of Applied Biosystems. Relative gene expression were analized in QuantStudio 5 software (ThermoFisher Scientific) and values were normalized to the expression of HPRT1 and beta-2 microglobulin (β2M) in the case of livers samples from young and old mice, whereas in the case of calorie restriction and glucose starvation treatments values were normalized to the expression of RNF219. Details of oligonucleotides are shown in supplementary table 4.

**Site-directed mutagenesis**

The different constructs harboring point mutations were cloned by site-directed mutagenesis based on the Quick-Change protocol (Stratagene). The PCR conditions were 25 cycles of denaturation at 95ºC for 60 seconds, annealing at 60ºC for 30 seconds, extension at 68ºC for 60 seconds per kilobase, final extension at 68ºC for 10 minutes. The parental (non-mutated) DNA was digested with 3 U of DpnI restriction enzyme for 1 hour at 37ºC. The positives clones selected were then verified by sequencing using an 3730 DNA Analyzer (Applied Biosystems, Foster city, CA, USA).

### Immunofluorescence

NIH3T3 cells were fixed with 4% paraformaldehyde for 7 minutes at room temperature, permeabilized for 5 minutes with buffer B (3% BSA, 0.2% Triton in PBS). After blocking in PBS containing 3% BSA in PBS for 1 hour, cells were incubated overnight at 4ºC with anti-Fibrillarin (Santa cruz, B-1, Sc-166001, 1:100) in Buffer B followed by 3 washes with 3% BSA-PBS and 30 minutes stained with fluorophore conjugated secondary antibody. Cells were stained with 1µg/ml of DAPI for 4 minutes and subsequently washed with PBS. Confocal images were taken with a confocal laser scanning microscopy (Leica, SP5) and digital images were analyzed using the Leica software (LAS AF lite).

### ChIP-seq and RNA-seq

For RNAseq, total RNA from 10 cm dishes containing WT and SirT7$^{-/-}$ exponentially growing MEFs was harvested using an RNEasy miniprep kit (Qiagen) according to the manufacturer instructions. Samples were sent to RUCDR, Infinite Biologics sequencing center (Rutgers University, NJ) for RNA-extraction, library preparation (Illumina Truseq chemistry), and sequencing (2x100bp PE reads, >60 x106 fragments per sample, Illumina HiSeq in Rapid mode). Reads were de-multiplexed and concatenated, and delivered as fastq files. Raw reads were assessed for quality with FastQC . Raw reads were trimmed with trimmomatic, aligned to the UCSC mm9 reference genome with HISAT2 (options: --dta), and converted to sorted and indexed BAM format with samtools. StringTie (in –eB mode) was used to estimate transcript abundance using the UCSC mm9 reference GTF file (obtained from Illumina iGenomes UCSC mm9 bundle). Differential expression was analyzed with DESeq2, allowing modeling terms for genotype, treatment, and batch.

Chromatin immunoprecipitation was conducted as described before[28]. Briefly, nuclear extracts from WT and SirT7$^{-/-}$ cells were first obtained and chromatin was prepared by sonication to an average fragment size of 300bp. For SirT7 ChIP, chromatin was diluted 8 times in Dilution Buffer and, for macro-H2A and H3K27K3m3 ChIP, chromatin was diluted 5 times. Antibodies were conjugated to beads and chromatin immunoprecipitation was conducted overnight at 4°C on a rotator. After washes and reversal of crosslinks, ChIP libraries were prepared using a ThruPlex DNA-seq library preparation Kit (Rubicon Genomics) following the manufacturer's protocol. ChIP libraries were then size selected with PippinPrep (Sage Science) for a final library fragment size of 350-400bp, and quantified with KAPPA qPCR. ChIP libraries were then sequenced on an Illumina HiSeq 2500 with 2x100bp or 2x75bp paired-end reads to a depth of >20x106 fragments per sample at the RUCDR, Infinite Biologics sequencing center (Rutgers University, NJ). Reads were de-multiplexed and concatenated, and delivered as fastq files.

**Analysis**

Raw read quality was first assessed with fastqc version 0.11.3, followed by adapter clipping and quality trimming with Trimmomatic version 0.36. Reads were then aligned to the mouse (UCSC mm9) reference genome with Bowtie with options –n 2 –k 10, and converted to BAM format with Samtools. ChIP, library, and alignment quality was evaluated computing library complexity (PBC), FRiP, insert size, and strand cross-correlation analysis. All samples passed the ENCODE quality standards for ChIP-seq studies. RPKM-normalized signal files were generated from BAM data using DeepTools version 2.4.2[55] using a bin size of 10bp.

SirT7 peaks were called in each biological replicate independently using MACS version 1.4.2 (--keep-dup auto --bw 450 --pvalue 0.05)[56]. Reproducibility of individual peaks was assessed with IDR (version 2.0.2) and the peaks passing a -log10 (global IDR) threshold of 1.3 across biological replicates were kept. SirT7-associated genes were called using the GREAT tool with IDR-passing peaks and the association rule of "Basal plus extension" with parameters Proximal=5.0 kb upstream, 1.0 kb downstream, plus Distal up to 10.0kb, including curated regulatory domains[34]. For macroH2A1.1 and H3K27me3 ChIP-seq datasets, we used ChromHMM to segment the genome into 4 learned states[57].

First bam files for ChIP and corresponding INPUT samples were binarized at a resolution of 200bp bins. Binarized data was submitted to the ChromHMM learnmodel program and allowed to learn 4 states. The learned model showed posterior enrichment for the following states: U1: no enrichment; U2: mH2A1 enrichment; U3: H3K27me3 enrichment; and U4: both mH2A1 and H3K27me3 enrichment. mH2A1-enriched regions were considered as the union of regions identified as states U2 and U4. mH2A1-enriched genes were called using the GREAT tool with mH2A1 regions as input with the association rule of “single nearest gene” within 10kb, including curated regulatory domains. Chromatin state transitions induced by GS treatment were computed relative to NT for each genotype. Briefly, we identified 200bp bins in GS with differing state calls than NT, and recorded the initial state (that occurring in the NT) and the final state (that occurring in the GS). We then took the subset of transition sites at the intersection of gene lists (+/- 3kb upstream and downstream of each gene). We quantified transition occupancy by counting the number of base pairs occupied by each transition type, and normalized this by the total number of base pairs that underwent any transition.  To ascertain the genomic context of these transitions, we submitted each transition type to the CEAS tool.

**Data availability**

All sequencing data for ChIP-seq and RNA-seq that support the findings of this study have been deposited in the National Center for Biotechnology Information Gene Expression Omnibus (GEO) and are accessible through the GEO Series accession number GSE128894. Proteomics data are available in Pride[58] with the dataset identifier PXD012020 and supplementary table 1.

### METHODS REFERENCES


38. You, W. *et al.* Structural Basis of Sirtuin 6 Activation by Synthetic Small Molecules. *Angew. Chem. Int. Ed. Engl.* 56, 1007–1011 (2017).
39. Sali, A. & Blundell, T. L. Comparative protein modelling by satisfaction of spatial restraints. *J. Mol. Biol.* 234, 779–815 (1993).
40. Berendsen, H., van der Spoel, D. & van Drunen, R. GROMACS: a message-passing parallel molecular dynamics implementation. *Computer Physics Communications* 91, 43–56 (1995).
41. Du, J. *et al.* Sirt5 is a NAD-dependent protein lysine demalonylase and desuccinylase. *Science* 334, 806–809 (2011).
42. Moniot, S., Schutkowski, M. & Steegborn, C. Crystal structure analysis of human Sirt2 and its ADP-ribose complex. *J. Struct. Biol.* 182, 136–143 (2013).
43. Davenport, A. M., Huber, F. M. & Hoelz, A. Structural and functional analysis of human SIRT1. *J. Mol. Biol.* 426, 526–541 (2014).
44. Gai, W., Li, H., Jiang, H., Long, Y. & Liu, D. Crystal structures of SIRT3 reveal that the α2-α3 loop and α3-helix affect the interaction with long-chain acyl lysine. *FEBS Lett.* 590, 3019–3028 (2016).
45. Pannek, M. *et al.* Crystal structures of the mitochondrial deacylase Sirtuin 4 reveal isoform-specific acyl recognition and regulation features. *Nat Commun* 8, 1513 (2017).
46. Luger, K., Rechsteiner, T. J. & Richmond, T. J. Expression and purification of recombinant histones and nucleosome reconstitution. *Methods Mol. Biol.* 119, 1–16 (1999).
47. Durkin, M. E., Qian, X., Popescu, N. C. & Lowy, D. R. Isolation of Mouse Embryo Fibroblasts. *Bio Protoc* 3, (2013).
48. Dignam, J. D., Lebovitz, R. M. & Roeder, R. G. Accurate transcription initiation by RNA polymerase II in a soluble extract from isolated mammalian nuclei. *Nucleic Acids Res* 11, 1475–1489 (1983).
49. Bilan, V., Leutert, M., Nanni, P., Panse, C. & Hottiger, M. O. Combining Higher-Energy Collision Dissociation and Electron-Transfer/Higher-Energy Collision Dissociation Fragmentation in a Product-Dependent Manner Confidently Assigns Proteomewide ADP-Ribose Acceptor Sites. *Anal. Chem.* 89, 1523–1530 (2017).
50. Perkins, D. N., Pappin, D. J., Creasy, D. M. & Cottrell, J. S. Probability-based protein identification by searching sequence databases using mass spectrometry data. *Electrophoresis* 20, 3551–3567 (1999).
51. Rosenthal, F., Nanni, P., Barkow-Oesterreicher, S. & Hottiger, M. O. Optimization of LTQ-Orbitrap Mass Spectrometer Parameters for the Identification of ADP-Ribosylation Sites. *J. Proteome Res.* 14, 4072–4079 (2015).
52. Zhang, J. *et al.* PEAKS DB: de novo sequencing assisted database search for sensitive and accurate peptide identification. *Mol. Cell Proteomics* 11, M111.010587 (2012).
53. Vakhrusheva, O. *et al.* Sirt7 increases stress resistance of cardiomyocytes and prevents apoptosis and inflammatory cardiomyopathy in mice. *Circ. Res.* 102, 703–710 (2008).
54. Worlein, J., Baker, K., Bloomsmith, M., Coleman, K. & Koban, T. The Eighth Edition of the Guide for the Care and Use of Laboratory Animals. 73, 98 (2011).
55. Ramírez, F. *et al.* deepTools2: a next generation web server for deep-sequencing data analysis. *Nucleic Acids Res.* 44, W160–5 (2016).

56. Zhang, Y. *et al.* Model-based analysis of ChIP-Seq (MACS). *Genome Biol.* 9, R137 (2008).
57. Ernst, J. & Kellis, M. ChromHMM: automating chromatin-state discovery and characterization. *Nature Publishing Group* 9, 215–216 (2012).
58. Vizcaíno, J. A. *et al.* 2016 update of the PRIDE database and its related tools. *Nucleic Acids Res.* 44, D447–56 (2016).
