## Supplemental Figures for "SirT7 auto-ADP-ribosylation regulates glucose starvation response through macroH2A1.1"

### SUPPLEMENTAL INFORMATION

**LIST OF SUPPLEMENTAL ITEMS**:

- **INCLUDED IN THIS FILE**:


Supplementary Figure 1. Related to Figure 1

Supplementary Figure 2. Related to Figure 2

Supplementary Figure 3. Related to Figure 4

Supplementary Figure 4. Related to Figure 5

Supplementary Figure 5. Related to Figure 5

Supplementary Figure 6. Related to Figure 5

Supplementary Figure 7. Related to Figure 6

Supplementary Tables S1-S4 Legends


- **NOT INCLUDED IN THIS FILE**:


Supplementary Tables S1-S4 (included separately)

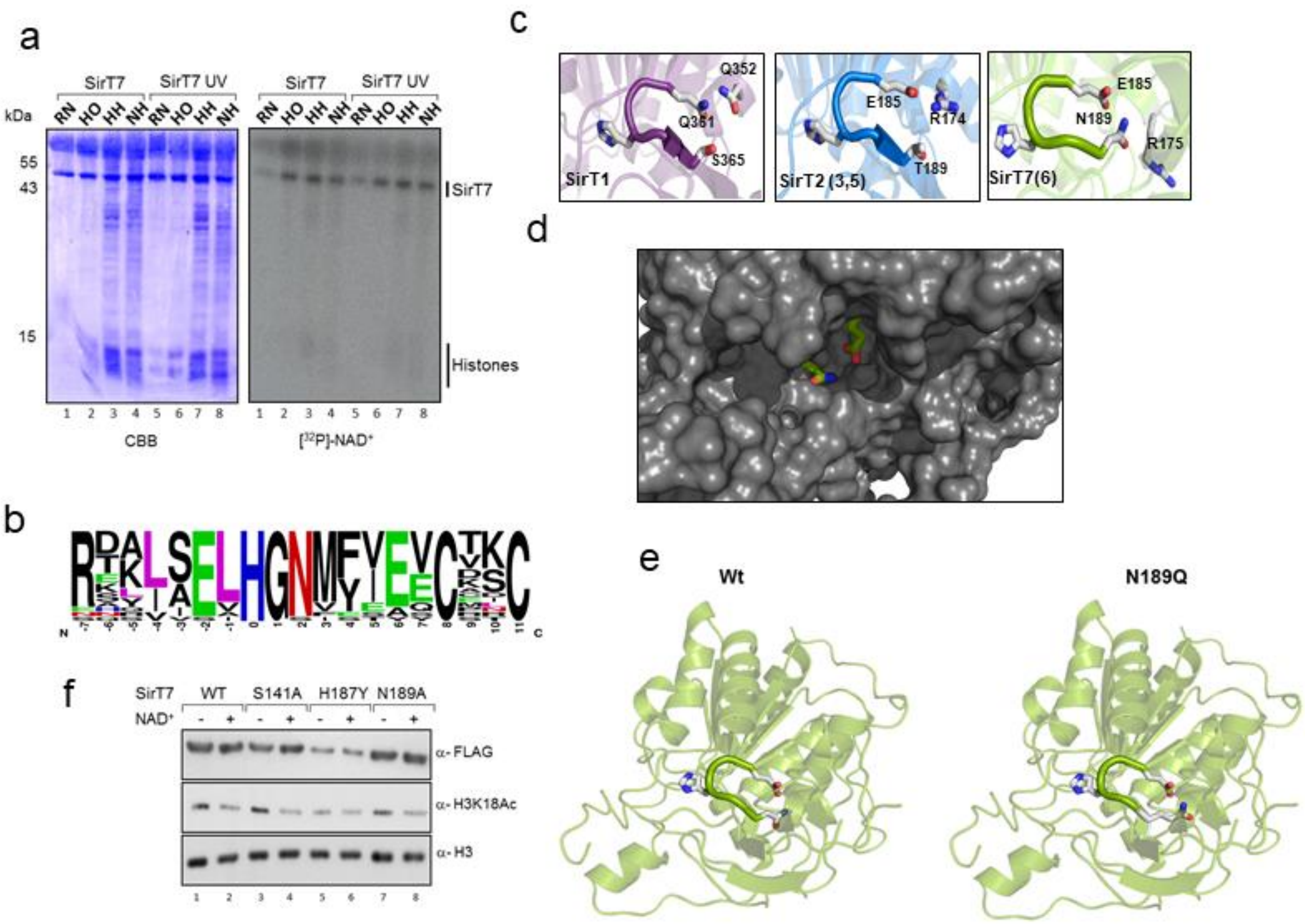


**Supplementary Fig.1**. **a,** ADPribosylation assay of purified SirT7 from untreated or UV-treated HEK293F cells incubated with the indicated histone substrates and [$^{32}$P]-NAD$^+$. RN:Reconstituted nucleosome; HO: recombinant histone octamers; HH: Hyperacetylated native core histones; NH: untreated native core histones. Coomassie blue-staining (CBB) and autoradiography are shown. **b,** Sequence-specific analysis of putative catalytic regions conserved in SirT6 and SirT7 orthologs as well other Sirtuins with known (SirT6 and SirT7 from humans to *C. elegans*; *P.falciparum* Sir2; *T. brucei* Sir2, *T. maritima* Sir2) or putative ADPRTs (SirT6 and SirT7 from *M. musculus*, *D. melanogaster*, *D. rerio*, *A. mississippiensis*, *A. mellifera*, *X. laevis*, and *G. gallus*). **c,** Comparative structure of the loop around H187 (in SirT7) between SIRT6/7 (right panel), SirT2/3/5 (middle panel) and SirT1 (left panel). While N189 is only present in SirT6/7, E185 is conserved in all of them except SirT1. In the case of SirT2/3/5 the glutamic residue is blocked by interaction with an arginine residue. **d,** Close view of the Structural model around the cavity, indicating the exposed position of the N189 and E185 residues. **e,** Structural molecular model of SirT7 WT (Left panel) and SirT7 N189Q (Right panel). The highly conservative amino acid substitution of N189 to glumatine (N189Q) within the loop around H187 and E185 does not alter the structure of the domain. Structural molecular models were based on the reported crystal structure of SirT6 (PDB 5M6F, 1.87 Å of resolution). **f,** SirT7-dependent deacetylation of H3K18ac. Purified Flag-tagged SirT7 (Wild Type, S141A. H187Y, and N189A) from HEK293F cells were incubated *in vitro* in the presence of [$^{32}$P]-NAD$^+$ and hyperacetylated histones. After the reaction proteins were subjected to SDS-PAGE followed by Western-blot with the indicated antibodies.

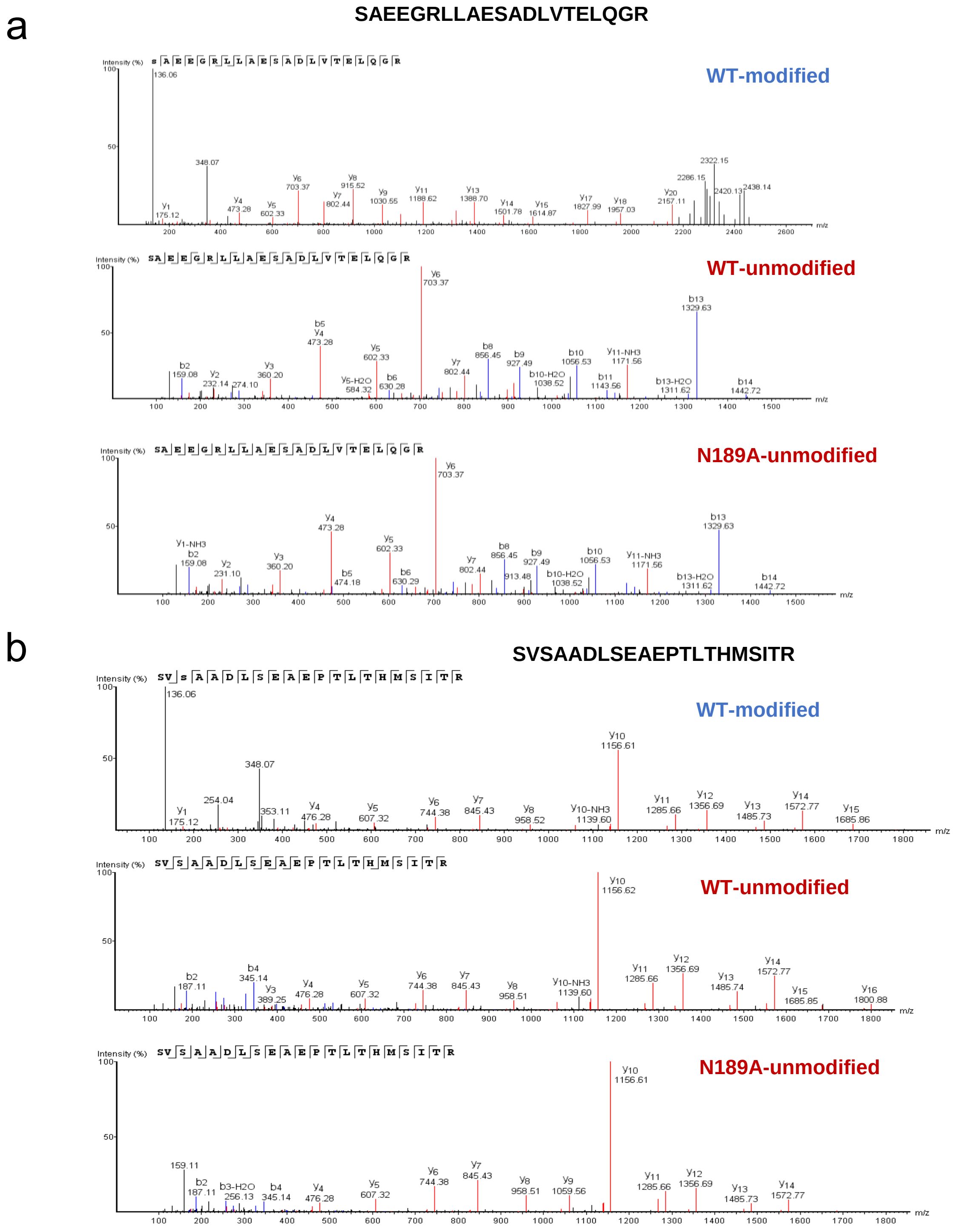


**Supplementary Fig.2**. **a-b,** Higher-energy collisional dissociation (HCD) spectrum of ADP-ribosylated and unmodified peptides identified in SirT7 wild type and SirT7 mutant (N189A). Although the exact localization site of the ADP-ribosylation could not be matched, the spectra corresponding to the WT samples show evidence of ADP-ribosylation with two of the most abundant ADP-ribose fragments (136-adenine and 348-adenosine mono-phosphate (AMP)).

a

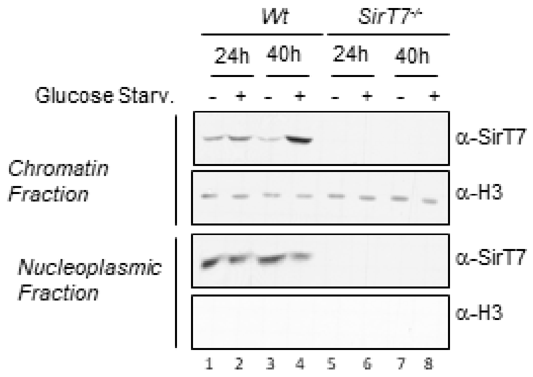


b

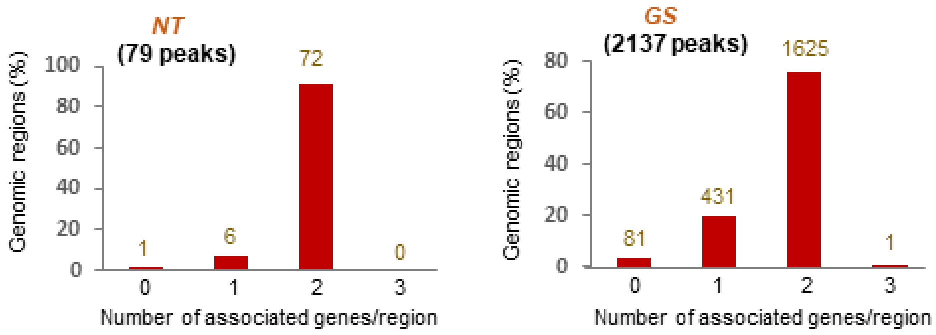


**Supplementary Fig.3**. **a,** Western blotting of endogenous SirT7 and histone H3 in chromatin and nucleoplasm fractions from *Wt* and *Sirt7*$^{-/-}$ MEFs cultured (+) or not (-) under glucose starvation. **b,** Comparison between the number of SirT7 associated genes per regions under normal cell growth (NT, left panel) or upon glucose starvation (GS, right panel) in MEFs cells by GREAT analysis (see Figure 4C-D and Methods). The analysis shows SirT7-binding sites assigned as putatively regulating each gene based on the following association rule: Proximal; 5.0 kb upstream, 1.0 kb downstream, plus Distal; up to 10.0 kb.

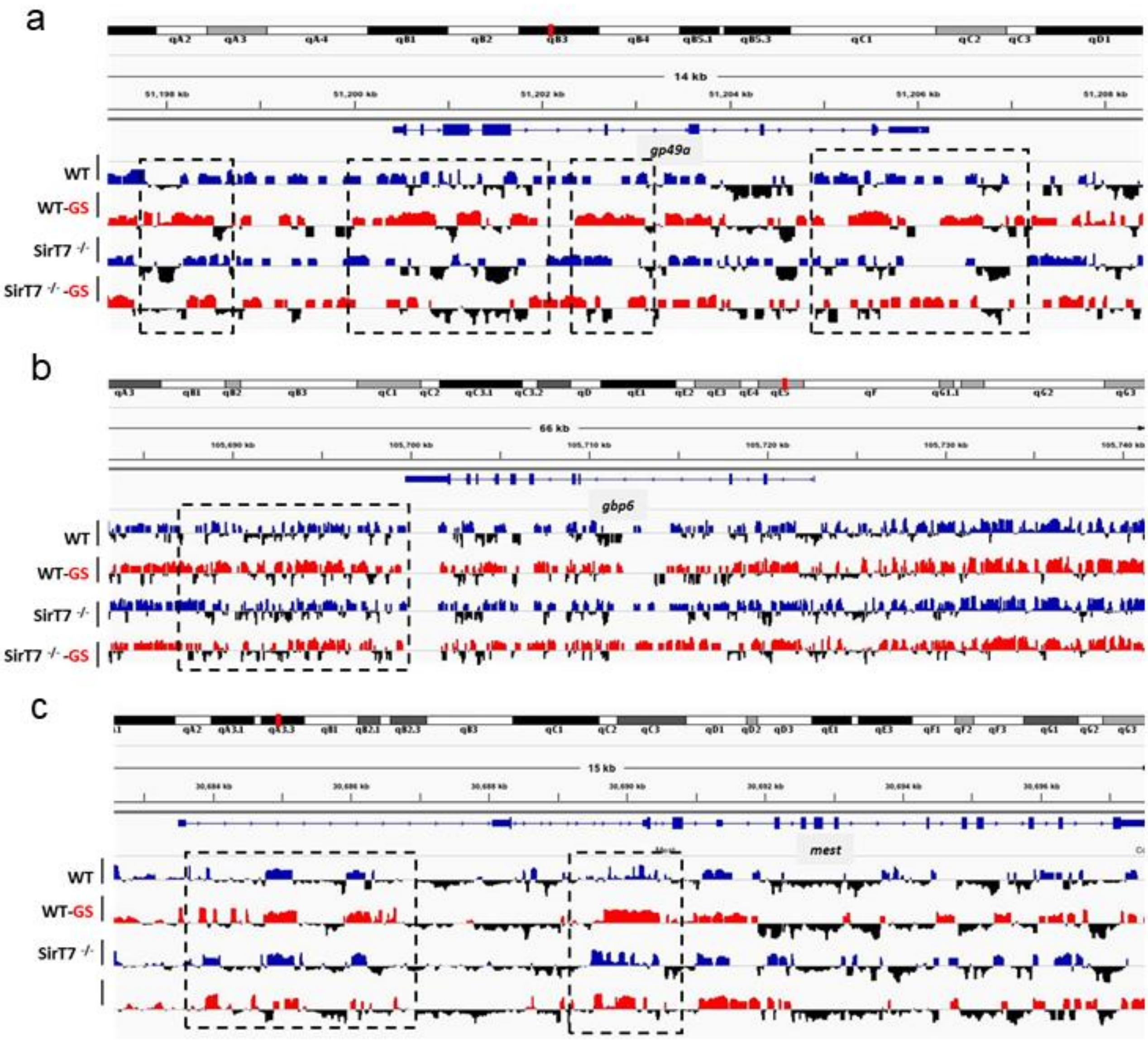


**Supplementary Fig.4**. **a-c,** Integrative Genome Viewer (IGV) examples of macroH2A1 ChIP-seq track peaks across genes and intergenic regions. Differentially macroH2A1-enriched regions upon GS (red for positive values) compare to non-treated (blue for positive values) wild type and *SirT7*$^{-/-}$ MEFs are highlighted in black dotted lines.

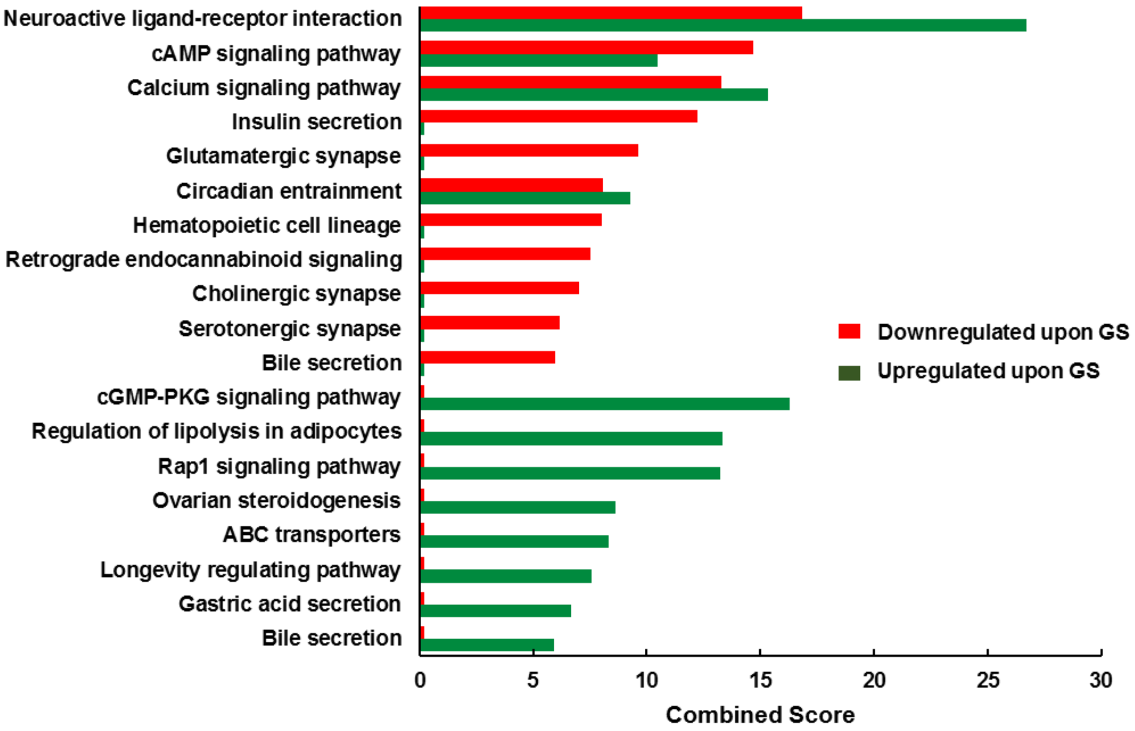


**Supplementary Fig.5.** KEGG´s cell signaling pathways affected upon GS compared to non-treated conditions (NT) in MEFs cells. The analysis was restricted to genes associated with SirT7 macroH2A1-enriched, and whose RNA expression was Down- and Up-regulated (277 and 143, respectively) upon GS. The different signaling pathways were rank-ordered by the combined score provided by enrich.

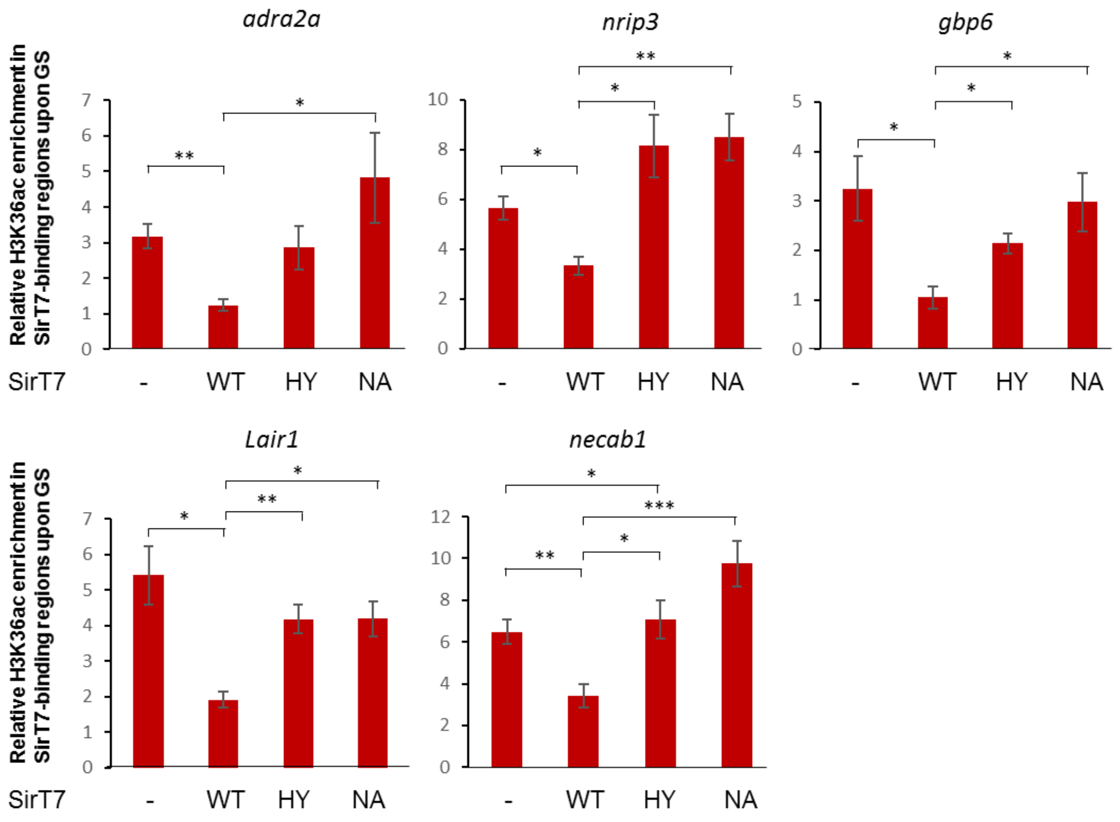


**Supplementary Fig.6**. ChIP analysis as in Fig.4c of H3K36ac enrichment upon GS in SirT7-binding sites associated to each gene. Quantification was generated from *n*=3 experiments, and *p*-values were calculated using two-tailed t-test (*: p<0.05; **: p<0.01; ***:p<0.005).

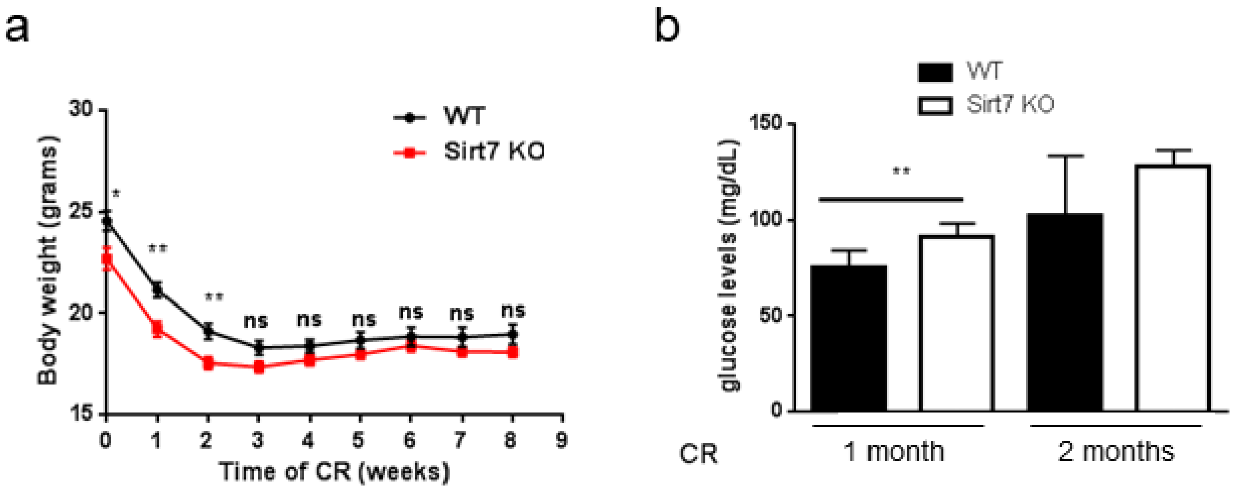


**Supplementary Fig.7. a,** Body weight monitoring at each indicated time point along calorie restriction (CR) treatment in wild type and *SirT7*$^{-/-}$ mice. **b,** Blood glucose levels during 1 month or 2 month of CR treatment in *Wt* and *SirT7*$^{-/-}$ mice. (*Wt* n=8; *SirT7*$^{-/-}$ n=6). p-values are calculated using two-tailed t-test (*: $p<0.05$; **: $p<0.01$).

### SUPPLEMENTAL TABLE LEGENDS

**Supplementary Table S1.** ADP-ribosylated peptides identified in SIRT7 WT and N189A analyzed using HCD and EthcD fragmentation methods. First data sheet includes ADP-ribosylated peptides of SirT7WT protein analyzed with Protein discover software. Second and Third data sheets show ADP-ribosylated peptides identified with PEAKS7 software in SirT7 WT and N189A mutant, respectively.

**Supplementary Table S2**. Mass spectrometry analysis of SirT7-HA elution. Protein identification of SirT7-associated protein by LC-MSMS analysis. The results were analyzed with MaxQuant software (1.6.2.6) using the built-in Andromeda search engine and the swissprot human database (March, 2018). The searches were filtered by a FDR at both peptide and protein level set to 1%.

**Supplementary Table S3.** Table with quantification of average chromatin mark enrichment (mH2A1 and H3K27me3), as log2(ChIP/INPUT), in promoter (-5kb to TSS) and gene body of genes associated with SirT7. Also included is RNA-seq Log2 Fold Change of NT vs GS in the WT and SirT7 KO and the distance between WT and KO Log2 Fold Change. Starting from all genes, we filtered to only include genes associated with SirT7 by GREAT analysis that are also positive for macroH2A1.1 enrichment. We then filtered to keep genes with an absolute log2 fold change in expression of > 0.6 in WT-NT versus WT-GS. Finally, we filtered to keep genes where the distance between the WT and KO expression log2 fold change with GS was > 0.3. The data is split between upregulated genes and downregulated genes based on the WT.

**Supplementary Table S4.** Oligonucleotides used in this study. Oligos for RT-qPCR, ChIP-qPCR and site-directed mutagenesis are included. All the oligonucleotides, with exception of mutagenesis with human SirT7, correspond to mouse sequences.
